# Mapping genetic risk mechanisms for immune-mediated diseases across human dendritic cell differentiation

**DOI:** 10.64898/2026.05.28.728586

**Authors:** Ofir Cohn, Chen Weng, Tianyi Ye, Anna-Lena Neehus, Chun-Jie Guo, Laila Barakat Norford, Emily Kao, Vijay G. Sankaran

## Abstract

Defining the cell types and mechanisms through which genetic variation operates is essential to understand the biological basis of disease. Although human dendritic cells (DCs) are crucial in regulating immunity, their rarity and limitations in available genomic data have hampered efforts to link inherited disease risk to specific DC subsets. Here, we present a single-cell multi-omic atlas of human DC differentiation from hematopoietic stem and progenitor cells (HSPCs) that includes chromatin accessibility and transcriptomic profiles to infer regulatory networks across DC subsets. By integrating this regulatory architecture with fine-mapped variants from hundreds of complex-trait genome-wide association studies, we systematically map inflammatory, autoimmune, and oncologic disease risk to DC subset-specific variant-to-gene mechanisms. For example, we identify a risk-associated variant that enhances the activity of a *PLD4* regulatory element in plasmacytoid DCs, thereby increasing the risk of developing systemic lupus erythematosus and other immune disorders. Collectively, these findings enable a deeper understanding of how complex immune diseases can emerge due to the impact of genetic variation acting in specific DC subsets.

## Introduction

Dendritic cells (DCs) originate from hematopoietic stem and progenitor cells (HSPCs) in the bone marrow and undergo several differentiation steps before maturing into different types of DCs, including conventional DCs (cDC1s and cDC2s) and plasmacytoid DCs (pDCs)^1^. DCs play a pivotal role in regulating immune responses while maintaining homeostasis by balancing immunity and tolerance. Through their functional plasticity, different DC subsets coordinate specific adaptive immune responses, such as by instructing various T helper cell populations to initiate their differentiation programs^2,3^. In multiple disorders, aberrant frequencies of circulating DCs are observed^4–7^. Furthermore, functional or developmental impairment of DCs may lead to a break in immune tolerance, underlying a wide spectrum of immunological abnormalities, including autoimmunity, allergies, impaired host defense, or dampened anti-tumor immunity^8–12^. DC dysregulation exerts a two-fold effect on disease pathogenesis: it directly alters the production of cytokines, such as type I interferons (IFNs), while simultaneously impairing antigen presentation capacity, thereby affecting T cell-mediated immunity. However, due to their infrequency, short half-life, and challenges in performing genetic modifications (without eliciting immune responses), most of our understanding of DC biology derives from mouse models, which show notable differences from humans^13–16^.

While genome-wide association studies (GWASs) have been successful in identifying disease-relevant inherited variation, knowledge of the underlying mechanisms remains limited in most cases, particularly for variants impacting extremely rare populations like cDC1s, transitional DCs (tDCs), and mature DCs (mDCs)^17^. A critical question for effective prioritization and functional validation of GWAS variants is through which gene, and particularly, in which cell type or cell state, the variants act. Many disease-associated variants reside within noncoding regulatory regions, where they influence gene expression by altering the epigenomic landscape (e.g., enhancer–promoter interactions). However, human DC studies that integrate genetic data with single-cell genomic data have been limited in scale and primarily focused on a single modality^18,19^. Although mouse DC studies have been useful, given the limited human data on these rare but important cell types, little progress has been made in identifying genetic variation associated with DC-specific disease mechanisms.

Here, we employ a tractable human DC differentiation system from HSPCs to deeply profile all DC subsets and their precursors, enabling single-cell multi-omic insights into these cells and their differentiation. Additionally, we compile *in vivo* single-cell transcriptome datasets from over 300 donors to create a comprehensive DC atlas to define the core transcriptomic signature across tissues and states. Our data provide key insights into the gene regulatory networks underlying human DC differentiation, which we validate in several cases. By leveraging this rich dataset in tandem with thousands of inherited variants associated with complex immune-mediated diseases, we generate a comprehensive map of complex diseases arising from altered regulation in specific human DC subsets. We link putative causal variants residing in regulatory regions to their potential causal genes in a cell-type-specific manner. Through these analyses, we identify unappreciated mechanisms for complex diseases. For instance, these insights lead to the identification and functional validation of inherited variation impacting phospholipase D family member 4 (*PLD4*) in pDCs that increases risk of systemic lupus erythematosus (SLE) and other inflammatory/autoimmune disorders. Together, we define a key and underappreciated role of human DCs in mediating the consequences of inherited genetic variation on a range of complex disorders, expanding the relevance of these rare, yet critical, immune cells in human disease.

## Results

### DC heterogeneity delineated by single-cell transcriptome analysis

Although DCs are central to the establishment of immunity and tolerance, the study of human DC biology has been partially challenged by the difficulty of capturing the full spectrum of DC lineages and differentiation states, particularly for rare DC subsets (**Fig. S1a,b**), and by the challenge in defining how developmental pathways and gene regulatory mechanisms are impacted by genetic variation in these cells^20,21^. To address these limitations, we implemented a tractable differentiation system^21,22^ that produces all characterized human DC subsets from cord blood-derived CD34^+^ HSPCs (**Fig. 1a**). Differentiated DCs were morphologically distinguishable by Wright-Giemsa staining and functionally validated by TLR agonist-induced DC maturation (**Fig. S1c,d**). To further characterize the culture-derived DCs and their precursors, we applied an integrated single-cell multiomics approach,Regulatory Multiomics with Deep Mitochondrial mutation profiling or ReDeeM (RNA-seq, ATAC-seq, and deep mitochondrial DNA mutation profiles)^23^, in over 25,000 Lineage^−^CD45^+^HLA-DR^+^ DC-enriched cells and precursors from three independent donors (**Fig. 1b**). Cluster analysis identified 24 cell clusters, which were manually annotated based on differential expression analysis and published signatures^18,24^. Importantly, DC subsets accounted for 93% of the cells assigned to cell clusters, such as cDC1 (*CLEC9A, CADM1*), pDC (*GZMB, CLEC4C*), cDC2 (*FCER1A,CLEC10A*), and mDC clusters (*LAMP3, BIRC3*) (**Fig. 1c-e**). The data showed high concordance across donors, as evidenced by shared ATAC-seq peak calls, gene expression clusters, and a positive correlation of motif activity (**Fig. S1eh**). We observed considerable diversity within the DC sub-populations. Specifically, we identified a cluster (annotated as ‘cDC2_6’) that had markers associated with tDCs (also referred to as AXL^+^SIGLEC6^+^ DCs (ASDCs))^25^ (**Fig. S1i**).

**Fig. 1.**
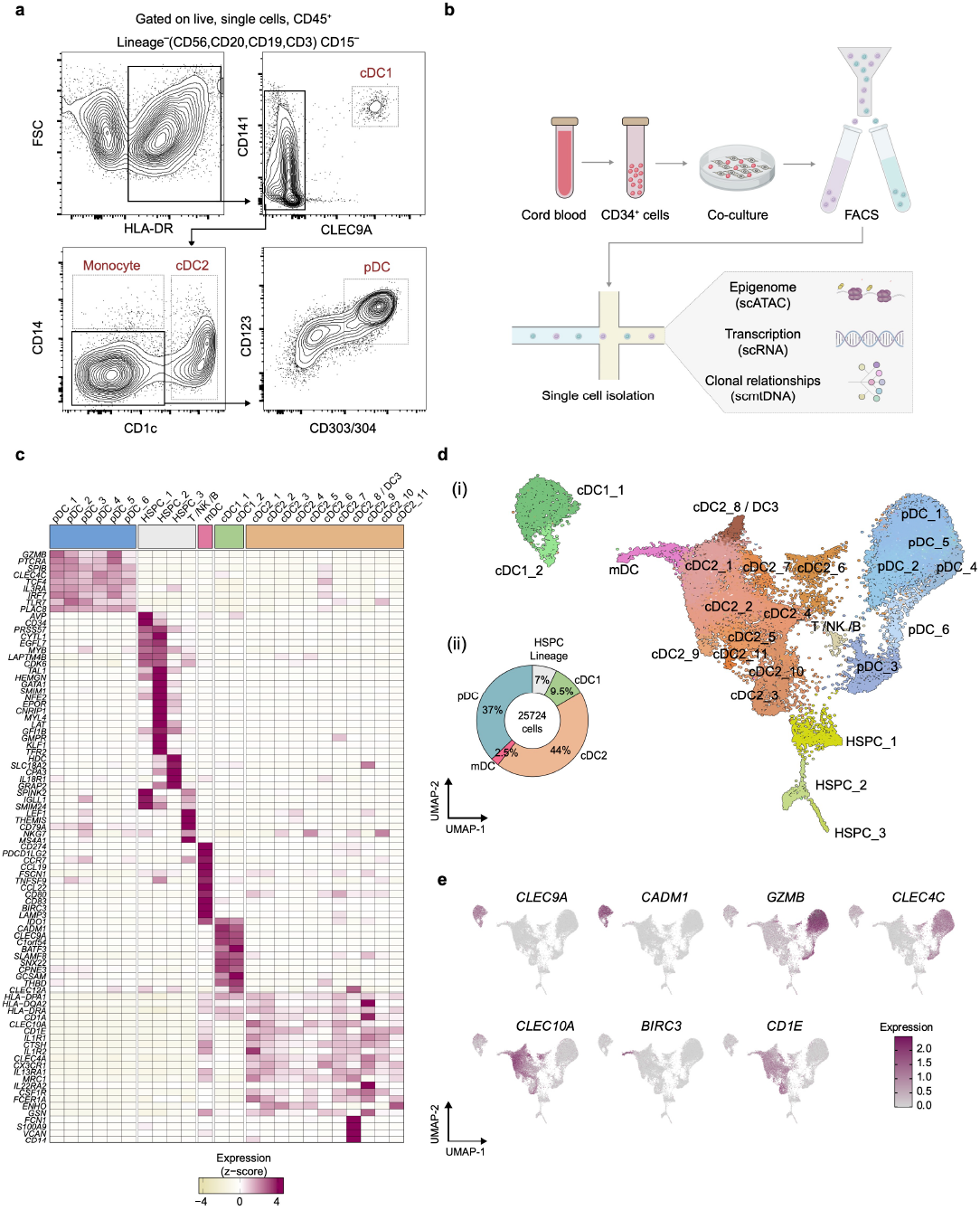
Single-cell transcriptomic profiling of DCs. **(a)** Representative flow plot of DC subsets after 15 d of co-culture of cord blood CD34^+^ HSPCs with mouse stromal cells (MS5). DC subsets were defined as Lineage^−^HLA-DR^+^ CD141^+^CLEC9A^+^ cDC1s, CD1c^+^ cDC2s, and CD123^+^CD303/304^+^ pDCs. **(b)** Experimental workflow. Lineage^−^CD45^+^HLA-DR^+^ cells (co-cultured as in **a**) were sorted and processed for scRNA-seq, scATAC-seq, and scmtDNA. **(c)** Heatmap showing the normalized average expression (*z*-scores) of selected marker genes for each cluster. **(d)** (i) Unsupervised dimensionality reduction of the data using Uniform Manifold Approximation and Projection (UMAP) embedding. The colors correspond to the identified clusters color-coded in the graphical legend. (ii) Donut plot showing the percentage of transcriptionally defined DC subpopulations and all other populations. **(e)** Average expression of DC markers projected on the UMAP plot. Abbreviations: NK, natural killer.

We also identified a distinct cluster (annotated as ‘cDC2_8 / DC3’) that exhibited the hallmark features of DC3s (such as *VCAN* and *FCN1*), while maintaining cDC2 transcriptomic signatures^26^ (**Fig. 1a** and **Fig. S1j**).

### A harmonized atlas of DC subsets reveals conserved gene programs

Establishing the relevance of our data from *in vitro* differentiated DCs to human disease requires assessing their epigenomic and transcriptomic resemblance to human primary cells from diverse tissues. Importantly, while DCs have multifaceted roles in regulating immune responses, it remains unclear how conserved their gene regulatory programs are across different tissues and disease states. In contrast to other leukocyte populations, DCs are rare under steady-state conditions in both blood and peripheral tissues, making high-resolution characterization challenging and limiting the ability to capture certain DC subsets and states^13,14,27^ (**Fig. S2a**).

Since nearly all tissue-resident DC subpopulations originate from a shared HSPC compartment, we sought to explore the core transcriptional programs defining DC subtypes. To do this, we first performed core signature alignment between our *in vitro* derived dataset and transcriptomic profiles of over 39,000 primary DCs from 13 independent single-cell RNA-seq datasets spanning multiple tissues and homeostatic/disease states (**Suppl. Table 1**). We observed strong conservation of DC subtype-specific transcriptional programs in the culture-derived DCs (**Fig. 2a**). In line with these findings, the chromatin accessibility landscapes were broadly conserved across tissues (**Fig. S2b**). Beyond the direct comparison to our data, an unbiased harmonization of DC subsets showed similarities in global transcriptome profiles, supporting the biological conservation of DC identities across a wide variety of physiological contexts (**Fig. 2b**). To establish a unified reference dataset, we integrated DC scRNA-seq datasets from 12 organs and from cultured DC subsets across 300 donors (**Suppl. Table 1**). Cell identities were assigned with high confidence using a logistic regression classifier trained at the low-hierarchy level, independent of tissue origin^28^. Notably, our *in vitro* derived dataset captures extremely rare subsets, such as cDC1s and mDCs, which are underrepresented in other datasets (**Fig. S2c,d**). Consistent with the matching scRNA-seq data, dimensionality reduction showed clear separation by DC subtypes rather than by study or tissue type (**Fig. 2c**). Analysis of the pan-tissue atlas confirmed the distinct expression of classic DC markers (e.g., *CLEC9A, CLEC10A*, and *GZMB*), but also highlighted several non-canonical conserved genes, including *MZB1* for pDCs, *IL7R* for mDCs, and *NCF2* for cDC2s (**Fig. 2d** and **Suppl. Table 2**). Beyond cross-tissue conservation, we identified localized gene expression programs associated with specific tissue environments. For instance, *CCL22* expression was more restricted to skin mDCs, whereas the *MX1* gene was predominantly expressed in pDCs from the peripheral blood of COVID-19 patients (**Fig. 2e,f** and **Fig. S2e,f**). Overall, our integrated DC atlas showed high concordance among DC-subtype transcriptomic signatures that are preserved across tissues in both healthy and disease states. In particular, the gene expression patterns and epigenomic profiles of primary tissue-resident DCs were closely mirrored by our *in vitro* culture system, supporting the use of our single-cell multi-omic dataset for identifying generalizable gene regulatory networks and mapping disease relevance within these networks.

**Fig. 2.**
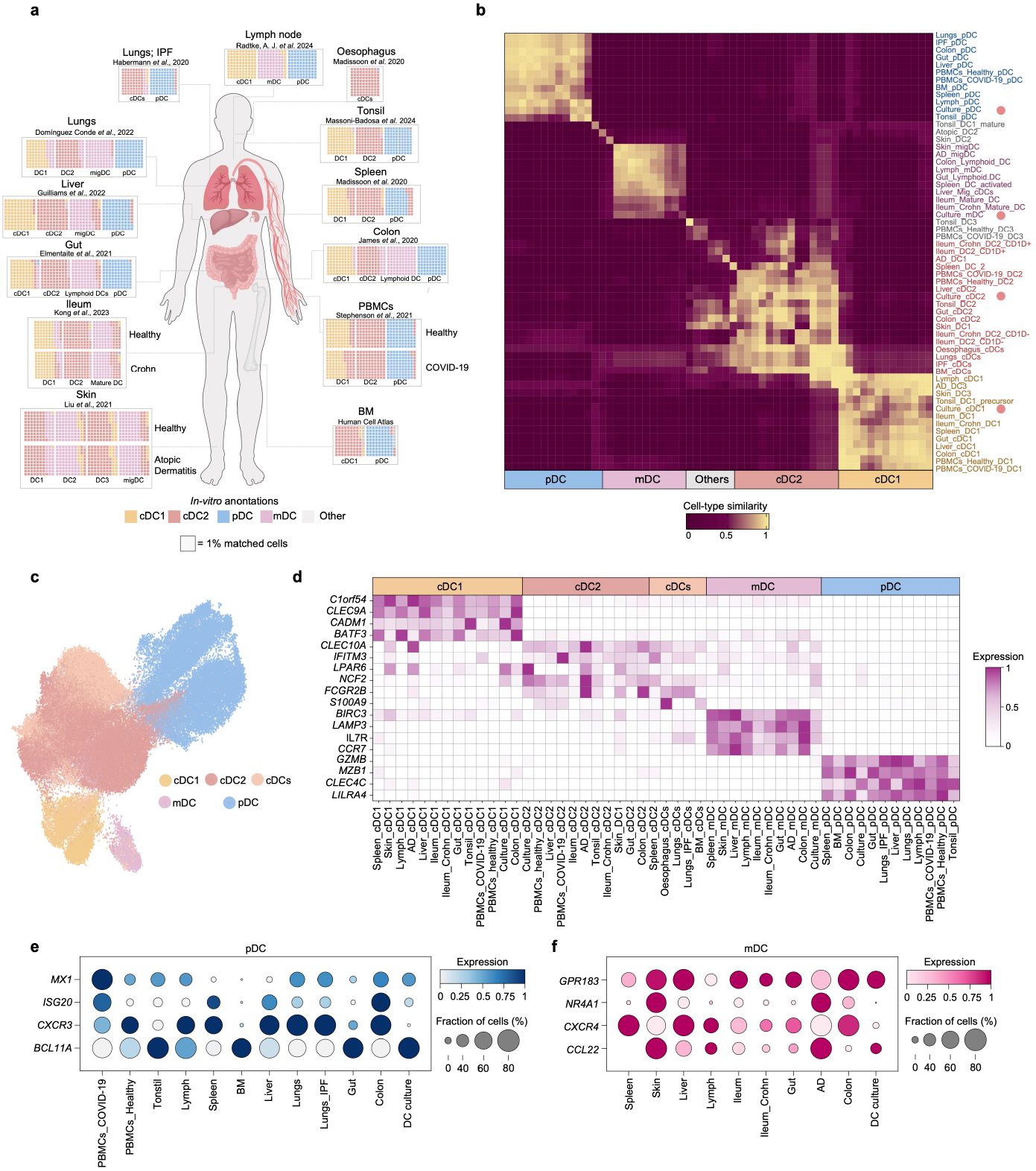
Conserved features of DC subtypes across tissues and states. **(a)** DC map demonstrating the subset similarities of different cell types in 12 organs. Cell type identities were determined by transferring labels from an *in vitro*-derived DC reference to the tissue query cells. Each plot represents a total of 100 units, where one square corresponds to 1% of the total query population for that organ. **(b)** Heatmap comparing DC subpopulations across datasets based on similarities inferred by CellHint. Cell types are color-coded by DC subtypes, with DC culture subsets highlighted. Color scale indicates CellHint similarity scores (0–1), where higher values denote greater transcriptional alignment between clusters across tissues. **(c)** UMAP visualization of cross-tissue integration of scRNA-seq data from DCs in health and disease. **(d)** Comparative expression of the top unique genes in DC subsets across different tissues. Color intensity represents the average expression level of each gene within a given cluster. **(e, f)** Dot plots showing the expression of representative tissue-specific genes across DC subpopulations and tissues of origin. Dot size indicates the percentage of cells expressing the indicated gene, and color intensity represents the average gene expression. Abbreviations: BM, bone marrow; AD, atopic dermatitis; IPF, idiopathic pulmonary fibrosis.

### Gene regulatory mechanisms underlying each DC-subset specification

Prior studies of human DCs have primarily examined transcriptional profiles^18,19^, while less is known about the underlying epigenetic programs and gene regulatory networks (GRNs) that govern DC differentiation and subtype-specific fate choices. Leveraging the paired multi-omic resolution of our *in vitro* data, we first analyzed chromatin accessibility profiles across the identified clusters (median 22,955 fragments per cell) to infer transcription factor (TF) activity. In agreement with the scRNA-seq profiles, DC subsets exhibited cell-type-specific TF motif enrichment patterns, including TCF4 in pDCs, BATF3 in cDC1s, and GATA2 in all the earlier hematopoietic progenitors (**Fig. 3a**). Dimensionality reduction and hierarchical clustering based on TF activity highlighted that pDCs have a distinct epigenetic landscape compared to other DC subtypes, suggesting that pDC specification is governed by a divergent regulatory architecture (**Fig. 3b** and **Fig. S2b**). To link these regulatory programs to DC differentiation dynamics, we used CytoTRACE2^29^, a deep learning framework that predicts the developmental potential of each cell. Of note, clusters ‘cDC1_2’, ‘cDC2_3’, and ‘pDC_3’ represented less differentiated states than their mature counterparts (**Fig. 3c** and **Fig. S3a,b**). Furthermore, this analysis revealed a transcriptional continuum of lineage-defining genes, transitioning from early progenitor markers (e.g., *KIT* and *MECOM*) to lineage commitment factors (such as *SPI1* and *ID2*), and finally to subset-specific determination (*BATF3, KLF4, TCF4*) (**Fig. 3d**). To systematically define the regulatory relationships that underlie DC development, we applied Pando^30^ to our data. Pando leverages multi-modal measurements (RNA and ATAC) to infer GRNs, identify sets of genes regulated by each TF module, and determine positive and negative regulatory interactions between TFs. Within this GRN inference, we incorporated RNA velocity information^31,32^ to map developmental trajectories onto the inferred differentiation graph. The global GRN uncovered distinct TF modules underlying early (including GATA1, MECOM, MEIS1) and late stages of DC development (including TCF4, BATF3, BCL6). Notably, GRN inference revealed a negative regulatory relationship between the early regulatory factor ZEB2 and the late factor ID2, consistent with the reported repressive role of ZEB2 in pre-DCs (**Fig. S3c**)^33,34^. To infer the regulatory circuits (i.e., active subnetworks of the GRN), we used differentially accessible regions enriched within each major DC subpopulation. This analysis recovered an ensemble of lineage-determining factors that control DC development (**Fig. 3e,f**). For example, this analysis highlighted PU.1 (SPI1), which is required for cDC (cDC1 and cDC2) formation^35^. In addition, we identified core components of the NF-κB complex (e.g.,NFKB1, REL, RELB) as central regulators of cDC maturation. This agrees with prior findings from mouse models and human clinical studies, which demonstrate that c-Rel is specifically essential for the production of IL-12 and IL-23 in activated cDC1s^11,36^. To obtain a complementary view of the regulatory programs across the DC lineage while accounting for non-linear TF–target relationships, we applied SCE-NIC+^37^. This approach identifies gene regulation modules (eRegulons) by integrating gene expression, chromatin accessibility, and motif enrichment to score their activity in individual cells. Dimensionality reduction based on eRegulon enrichment scores revealed a clear separation across DC lineages (**Fig. S3d,e**), consistent with the regulatory structure inferred from the GRN analysis. Taken together, by defining the key TFs that underpin human DC specification and maturation, we provide a comprehensive analysis of the DC regulatory landscape.

**Fig 3.**
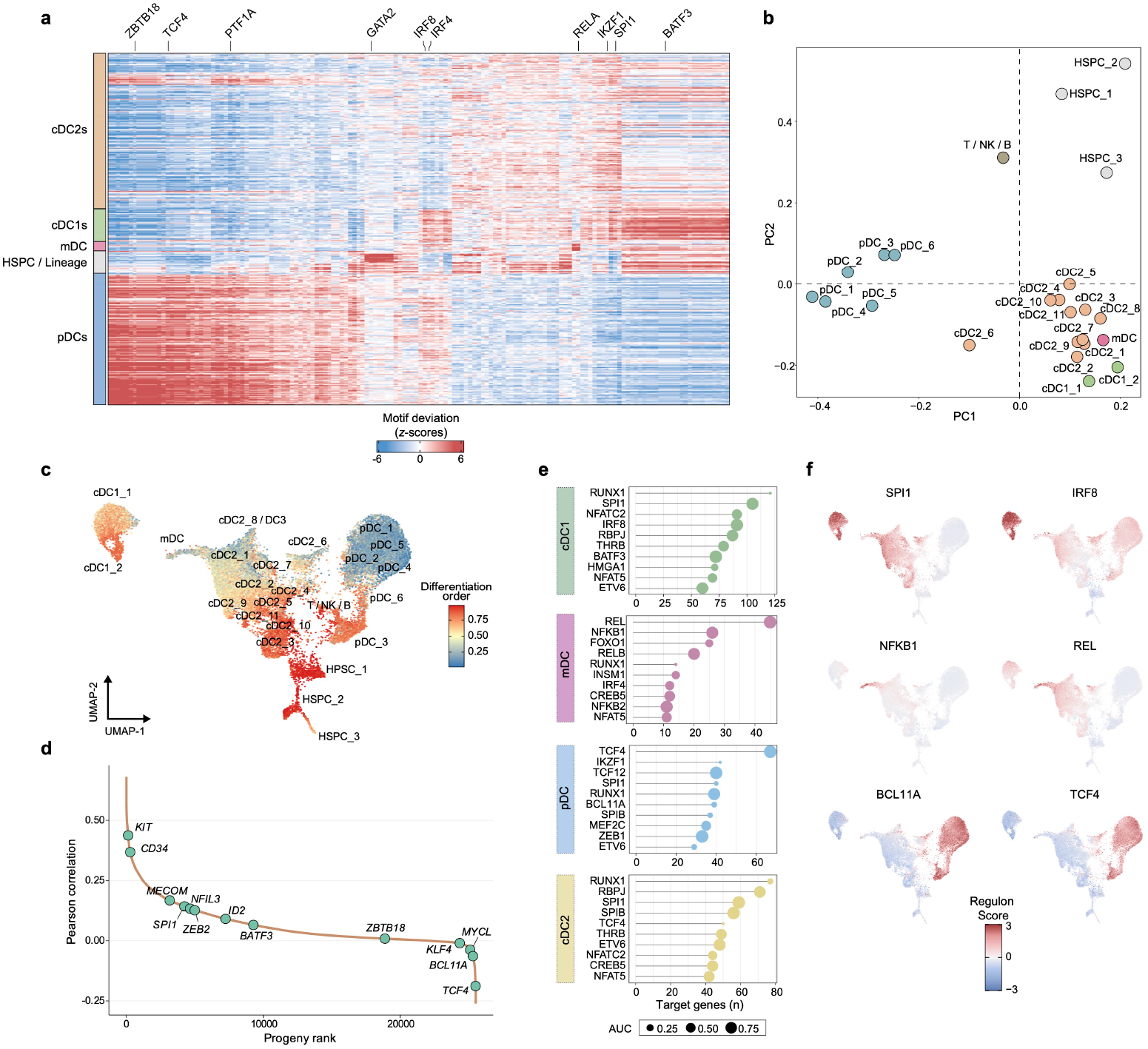
Gene regulatory networks of DC subtypes during differentiation. **(a)** Heatmap representation of chromVAR bias-corrected deviations for the top 200 variable TFs across all cell types. **(b)** PCA based on the TF activity scores calculated by chromVAR in **a. (c)** UMAP of clusters colored by developmental potency. **(d)** Gene expression–CytoTRACE2 progeny score Pearson correlations. Selected genes are highlighted in green and ranked by their correlation coefficient. **(e)** Lollipop plot showing the top ten TFs ranked by their number of connections. Node size represents the area under the curve (AUC) values of chromVAR bias-corrected deviations across groups. **(f)** UMAP projection of cell clusters colored by module score for each TF regulon in individual cells.

### Alterations in DC subset gene programs and developmental trajectories

To elucidate how DC-subset-specific GRNs are organized along differentiation trajectories, we first mapped the transition from precursor to terminal states across the major DC lineages (**Fig. 3c** and **Fig. S3a,b**). By correlating TF expression and chromatin accessibility with RNA velocity, we identified trajectory-specific regulatory networks. For example, *GATA2* and *ZEB2* displayed dynamic expression patterns along the cDC1 pseudotime trajectory, with high expression at early stages followed by progressive repression at the later stages. In contrast, factors such as *TCF4* and *IRF8* exhibited sustained expression with a gradual increase along the trajectory path toward pDCs (**Fig. 4a,b**). By incorporating our pseudotime trajectories and SCENIC+ analyses, we extracted key positive regulators of pDC fate and the corresponding negative regulators of cDC1 commitment. This further substantiated cell-type-specific regulatory activity for ZEB2 and TCF4 and highlighted additional factors such as BCL11A^38^ and ETS1. *ETS1* is highly expressed in pDCs and might be involved in the regulation of interferon-inducible genes produced by pDCs (*IFIT1, IFI44*, and *EIF2AK2*)^39^ (**Fig. 4c** and **Fig. S4a,b**).

**Fig 4.**
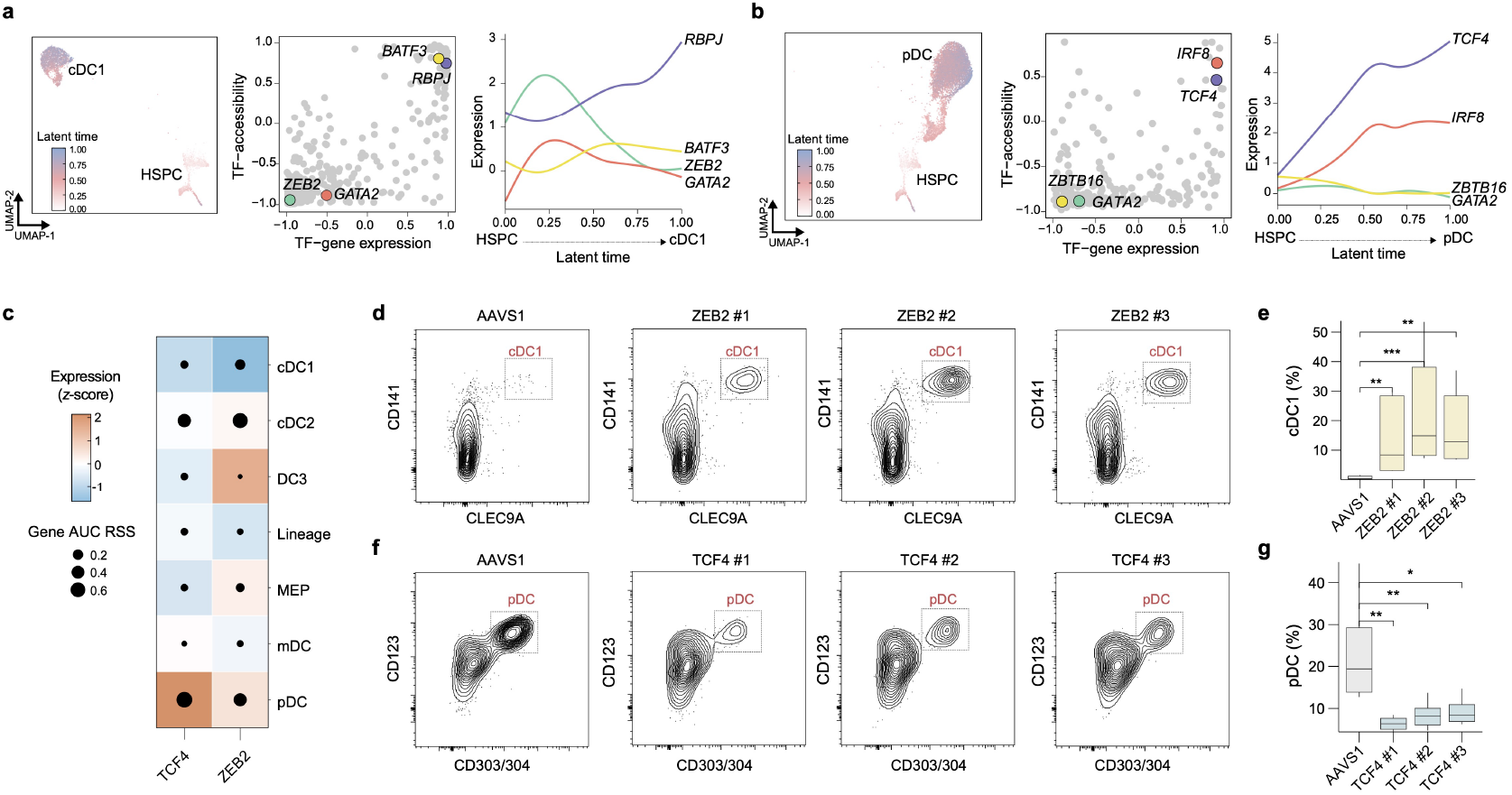
Regulatory mechanisms in DC subset differentiation. **(a-b)** Pseudotemporal cell type progression from HSPC to cDC1 (**a**) or pDC (**b**). Middle: Gene expression-ATAC correlations with latent time as inferred by scVelo for cDC1 (**a**) or pDC (**b**). Right: Pseudotemporal gene expression changes during DC development for each group. **(c)** Heatmap showing TF expression of the eRegulon on a color scale and cell-type specificity (RSS) of the eRegulon on a size scale as inferred by SCENIC+. **(d-g)** Representative flow cytometry analysis (d, f) and quantification of the proportions (e, g) of the indicated DC subsets following CRISPR-Cas9-mediated knockout of *ZEB2, TCF4*, or *AAVS1* (control). Statistical analysis: Mann–Whitney test (*n* = 8 donors). \**P* < 0.05; \*\**P* < 0.01; \*\*\**P* < 0.001.

Although genetic perturbations in mouse models have been commonly employed and provided foundational insights into DC regulation, they may not reflect the developmental pathways of human DCs^16,40,41^. Given that genome editing of *ex vivo*-isolated human DCs is often limited by unintended immune activation or restricted to mature developmental stages, we reasoned that our HSPC-derived DC platform could be exploited for genetic perturbations at the progenitor level. To directly validate our trajectory inference, we performed CRISPR-Cas9-mediated deletion of key regulatory nodes during DC differentiation from HSPCs. Consistent with our model, deletion of the pDC positive regulator *TCF4* reduced pDC fractions, while perturbing cDC1 negative regulator *ZEB2* dramatically increased cDC1 abundance (**Fig. 4d-g**). While cDC subsets remain unaffected in *Tcf4* knockout mice^42,43^, *TCF4* downregulation in our human system led to a moderate increase in cDC1 output, supporting the conserved antagonism between *Tcf4* and *Id2* during DC development (**Fig. S4c-e**)^44,45^. In line with these findings, when leveraging the deep mtDNA profiles obtained serendipitously from the ReDeeM analysis and by applying MitoDrift^46^ to infer clonal relationships, pDCs appeared to have closer clonal relationships with cDC1s in comparison to cDC2s (**Fig. S4f,g)**. This aligns with the close developmental relationships observed in clonal tracing of *Cx3cr1*^+^ DC progenitors in mice^47^ and in the relationships inferred through our GRN reconstruction. In summary, the *in vitro* DC differentiation platform from HSPCs provides a tractable human system to causally link gene regulatory programs to DC fate decisions.

### Identifying disease-critical cell types

Having established the regulatory architecture governing human DC specification, we next investigated how inherited genetic variation shapes these programs and contributes to the variable immune responses observed across multiple disorders. While GWASs have identified thousands of disease-associated loci, linking these variants to the relevant immune cell types and states remains a major challenge, particularly for rare cell populations and subtypes. To identify DC subsets and differentiation states that underlie genetic risk for complex immune-mediated diseases, we integrated fine-mapped variants identified from 696 complex-trait GWASs^48^ with our single-cell epigenomic profiles using a network propagation framework, implemented in SCAVENGE^49^ (**Fig. S5a**). SCAVENGE overlaps posterior probabilities from statistical fine-mapping with single-cell epigenetic profiles and calculates the enrichment strength of disease-associated variants by trait-relevance scores (TRSs) for each cell. Using this approach, we identified novel cell-type-specific associations between DC subpopulations and immune-related disorders^50–56^ (**Fig. 5a**). For instance, we observed significant enrichments of DC subpopulations in systemic autoimmune and inflammatory skin disorders (e.g., rheumatoid arthritis (RA), vitiligo, atopic dermatitis), indicating the critical role of DC subtypes in shaping tissue-specific immune responses. For cDC2s, we detected diverse disease associations, aligning with their established transcriptional and phenotypic heterogeneity^57,58^. In this regard, cluster ‘cDC2_6’, which expresses tDC markers (**Fig. S1i**), exhibited similar associations to those of pDCs. Despite overall similarities, clusters in earlier differentiation states (e.g., preDCs) displayed subtle differences in the enrichment of variants associated with certain phenotypes, suggesting differentiation stage-selective variation in chromatin accessibility. For example, ‘cDC2_3’, which represents a pre-cDC2 state (**Fig. S3a,b**), showed a weaker association with atopic dermatitis than the more mature ‘cDC2_1’ cluster. We then performed permutation testing for each cell to determine enrichment for a given trait. As shown in low-dimensional uniform manifold approximation and projection (UMAP) space, cDC1s and cDC2s were enriched with genetic variants predisposing to seborrheic keratosis and atopic dermatitis, respectively, while pDCs were enriched for risk of soft tissue infections (**Fig. 5b,c**).

**Fig 5.**
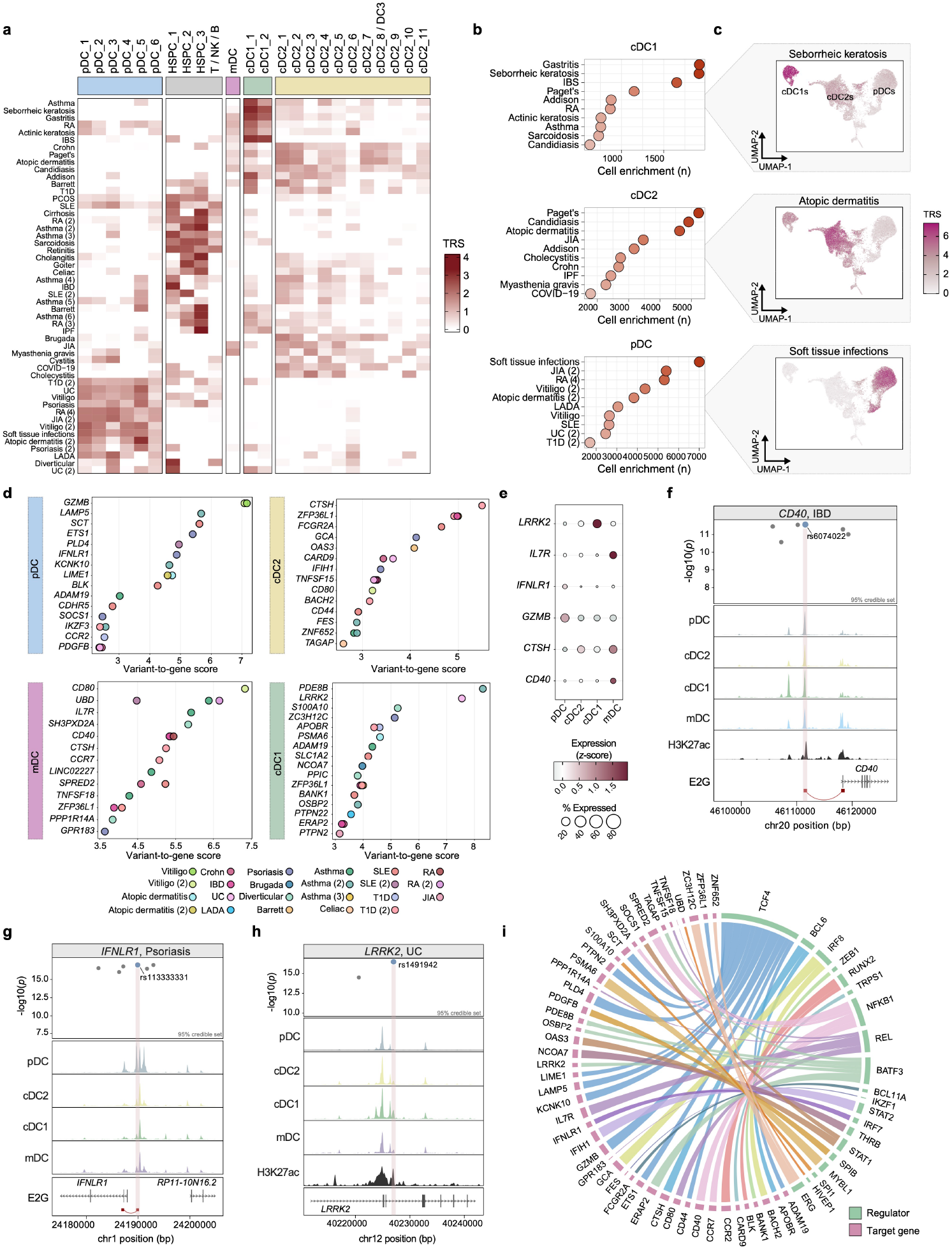
Relationship of DC subsets to diseases. **(a)** Heatmap showing the mean trait relevance score (TRS) (z-scores) of indicated traits for each cell type. **(b)** For the indicated trait, the number of enriched cells was defined by using network propagation scores yielded from SCAVENGE analysis (P value < 0.05, 1,000 permutations). **(c)** UMAP of indicated trait, colored by the SCAVENGE TRS. **(d)** Dot plots of scE2G-linked target genes by cell type. Each dot represents a target gene colored by associated trait and ranked by a combined scE2G score (see Methods).**(e)** Dot plot of the indicated genes across cell types. Dot size indicates the percentage of cells expressing the gene, and color intensity represents the average gene expression. **(f-h)** Normalized scATAC-seq signal coverage tracks showing cell-type-specific chromatin accessibility alongside GWAS summary statistics for the indicated trait, with a zoomed-in view of fine-mapped variants at the locus. Candidate enhancer regions are highlighted, with corresponding H3K27ac ChIP-seq signal. **(i)** Circular chord diagram showing relationships between TFs (green) and target genes (red), as inferred by SCENIC+ triplet rank. The thickness of the lines corresponds to the z-scored (inverted) triplet rank. Abbreviations: IBS, irritable bowel syndrome; T1D, type 1 diabetes; PCOS, polycystic ovary syndrome; IBD, inflammatory bowel disease; IPF, idiopathic pulmonary fibrosis; JIA, juvenile idiopathic arthritis; UC, ulcerative colitis; LADA, latent autoimmune diabetes in adults.

Given that complex diseases are typically driven by the coordinated activity of multiple cell types across tissues, we reasoned that disease-associated signals identified by SCAVENGE would not be universally DC-specific. To explicitly assess specificity, we expanded our analysis to include non-dendritic immune populations derived from PBMCs, such as monocytes, T cells, and B cells. This analysis identified disease–cell-type associations that were preferentially enriched outside the DC compartment, such as Paget’s disease in monocytes^59^ and primary biliary cholangitis in T and B cells^60,61^. However, this also highlighted several instances of DC-specific involvement in the pathogenesis of traits such as atopic dermatitis^62^ and sarcoidosis^63^ (**Fig. S5b,c**). To further assess the robustness of our findings, we compared the genetic associations derived from single-cell results with those obtained from pseudobulk populations. In addition to well-established phenotype/cell type associations, such as pDC enrichment in SLE, our analysis revealed previously underexplored associations, including enrichment of pDCs in juvenile idiopathic arthritis (JIA) and RA (**Fig. S5d-f**).

Our genetic linkage supports the recent findings regarding the pathogenic role of pDCs within the inflamed joint. First, beyond their canonical role in type I IFN production, pDC-derived TNF has emerged as a driver of RA pathogenesis^64,65^. Second, pDCs are considerably more abundant in patients with JIA^66,67^. Next, we expanded our survey to hundreds of diseases from the FinnGen Consortium^68^. This analysis revealed distinct, but frequently overlapping, connections between specific DC subpopulations and various phenotypes, including many for which the phenotype-driving cell types have yet to be defined (**Fig. S5g-i** and **Suppl. Table 3**). Consistent with our previous skin associations, we observed a strong enrichment of DC-specific genetic risk for cDC2s with atopic dermatitis, while skin infections were enriched in pDCs, as in the case of our independent analyses discussed above. These analyses indicate that genetic risk for inflammatory and autoimmune diseases is not distributed uniformly across the DC compartment but preferentially associates with specific DC subpopulations.

Finally, despite the critical role of DCs in cancer immunity^69^ and the promising development of DC-targeted therapies^70^, our understanding of the cell-type-specific genetic drivers of cancer susceptibility remains limited, particularly for pDCs, whose role in cancer is not well established. To this end, we explored the relevance of DCs across GWAS data spanning 34 malignancies representing 14 major cancer types. We found that cDCs were significantly enriched for genetic variants predisposing to rectal and pancreatic cancers, while pDCs were enriched for breast cancer risk, consistent with their role in facilitating tumor growth through immunosuppression^71^ (**Fig. S5j**).

### Variant-to-gene mapping reveals subtype-specific disease risk targets

Beyond cell-type associations, we sought to explore the regulatory mechanisms underlying these signals and delineate target genes. Here, we used the single-cell enhancer-to-gene(scE2G)^72^ framework to construct cell-type–specific variant–gene maps (**Fig. 5d** and **Suppl. Table 4**). We identified several key genes involved in DC functions, such as maturation and migration (**Fig. 5e**). For instance, the RA/inflammatory bowel disease (IBD) risk variant rs6074022^73^ was linked to an enhancer targeting *CD40* specifically in mDCs (**Fig. 5f**). The CD40-CD40L axis plays a critical role during the immune response by enhancing DC activation, including the upregulation of co-stimulatory molecules and pro-inflammatory cytokines. Consequently, constitutive *CD40* expression might drive inappropriate activation of autoreactive T cells, inducing Th1 and Th17 cytokine responses that directly result in tissue damage and intestinal inflammation^74,75^. We also observed a genetic convergence between asthma susceptibility and *IL7R* expression in mDCs^76^ (**Suppl. Table 4**). In DCs, IL7R is part of the thymic stromal lymphopoietin (TSLP) receptor complex and thus facilitates TSLP-driven DC activation to initiate allergic Th2-mediated immune responses^77–79^. Notably, a highly protective type 1 diabetes (T1D) missense variant (rs2289702, OR = 0.77)^80–82^ was linked to Cathepsin H (CTSH) within the cDC2 and mDC subsets (**Fig. S5k**). Cathepsins are central to DC function, acting as critical endolysosomal proteases required for the proteolytic cleavage and activation of Toll-like receptors (TLRs)^83^. Given that viral infections, particularly enteroviruses, are implicated as environmental triggers for T1D^84^,and that the rs2289702 allele provides protection in early childhood (<7 years)^80^, this regulatory linkage suggests that genetically reduced *CTSH* expression likely blunts TLR-mediated recognition of viral infections in cDC2s, thereby preventing the loss of tolerance in the periphery and the onset of T1D. Similarly, in pDCs, our analysis linked psoriasis-associated variants to *IFNLR1* (**Fig. 5g**), which encodes the receptor subunit for Type III interferons (IFN-III) (also known as IFN-λ). Given the reported infiltration of pDCs into lesional skin and the high local IFN-λ1 expression^85,86^, the identification of a protective variant (rs113333331, OR =0.80)^87^ linked to an enhancer regulating *IFNLR1*, might reduce pDC responsiveness to tissue inflammatory signals. Furthermore, our analysis linked several vitiligo-associated risk variants (e.g., rs2273844)^88^ to *GZMB* (granzyme B) specifically in pDCs (**Fig. S5l**). *GZMB* is a hallmark of the pDC transcriptional signature (**Fig. 1c)**. However, its precise role in autoimmunity has remained unclear. Previous studies have demonstrated that pDCs deliver GZMB to T cells to actively suppress their proliferation^89,90^. Because the destruction of melanocytes in vitiligo is driven by overactive T cells^91^, this regulatory linkage suggests that genetic risk at this locus impairs the pDC-mediated delivery of GZMB to T cells. Finally, we identified the *LRRK2* (leucine-rich repeat kinase 2) locus as a target of the risk variant rs1491942^73^ in ulcerative colitis (UC) (**Fig. 5h**), acting specifically within the cDC1 compartment. While *LRRK2* polymorphisms are classically studied in the context of Parkinson’s disease (PD)^92,93^, emerging evidence strongly implicates LRRK2 in peripheral immune responses and gut inflammation^94,95^. Consistent with this, we found that *LRRK2* is highly expressed in cDC1s (**Fig. 5e**) and might act as a critical regulator of DC migration via the modulation of intracellular calcium signaling^96^.

Having mapped these individual risk variants to their target genes, we next leveraged our GRN map to identify the core TFs governing these phenotype-associated loci. For the majority of pDC-specific regulatory elements and target genes, such as *GZMB* and *PLD4*, we identified the lineage-defining TF TCF4 as the primary putative regulator. mDC genes, including *CD40, CD80*, and *IL7R*, appeared to be largely governed by NF-κB family members, underscoring that these genetic risk variants might act within the core machinery of DC activation and inflammatory signaling (**Fig. 5i**).

In summary, our data demonstrate that autoimmune/inflammatory risk variants do not act uniformly in the DC compartments and might exploit the unique biological characteristics and regulatory mechanisms specific to each subset, thereby mediating different potential pathogenic mechanisms.

### pDC-specific *PLD4* variation in SLE

Given the strong enrichment of inherited genetic risk for SLE in pDCs (**Figs. 5a** and **6a**), we sought to identify specific genetic variants and regulatory mechanisms through which this variation might perturb pDC function. To determine whether this genetic predisposition is accompanied by measurable alterations in pDC abundance *in vivo*, we analyzed DC subset composition in scRNA-seq data from SLE patients and healthy individuals^97^. Based on known markers (e.g., *CLEC9A, LILRA4, FCER1A*), we identified distinct DC populations with transcriptomic resemblance to our culture system (**Fig. S6a-c**). Given the reported alterations in circulating pDC numbers in SLE patients^98,99^, we examined the differential composition of DC subsets in SLE patients and healthy individuals using two complementary statistical frameworks, Milo^100^ and Sccomp^101^. Milo was used to identify cell states enriched or depleted independently of discrete clustering, while Sccomp provided a robust model for compositional shifts across defined cell types. Across both methods, we found significant and concordant differences in DC subtype abundance. In particular, SLE samples demonstrated a significant decrease in circulating pDC proportions (**Fig. 6b** and **Fig. S6d**), which aligns with previous reports of pDC infiltration in target organs involved in immune attack (e.g., kidney)^102,103^. To functionally validate the candidate genetic regulators underlying SLE risk in pDCs, we first prioritized potential drivers by integrating our paired scRNA-seq and scATAC-seq data. Using the scE2G framework, we mapped SLE-associated variants to target genes. Among these regulatory elements, we identified four variants linked to *PLD4*, which also ranked as one of the top target genes (**Figs. 5g** and **6c**). PLD4 is a 5′ lysosomal exonuclease that generates the ligands for TLR7 to modulate IFN production^104,105^. Indeed, *PLD4* deficiency leads to a variety of autoimmune phenotypes, including the development of lupuslike features^106–108^. Yet, whether and how common variants alter PLD4 function and thereby impact autoimmunity is unknown. Furthermore, in contrast to its paralog *PLD3*, which is broadly expressed in other immune cells^105^, *PLD4* was exclusively expressed by pDCs and significantly upregulated in SLE patients (**Fig. 6d** and **Fig. S6e**). Among the four *PLD4*-linked variants, we further prioritized the variants by integrating pDC-specific ATAC-seq and H3K27ac profiles to identify active chromatin regions, which pinpointed rs2841277 as the most likely causal variant at this locus (**Fig. 6e,f**). In addition to SLE, analysis of independent GWAS datasets revealed associations between the *PLD4* variant rs2841277 and a broad spectrum of other autoimmune disorders, including RA, Sjögren’s syndrome, and scleroderma that showed the same effect direction as for SLE (**Fig. 6g**). To evaluate the cell-type-specific effects of rs2841277 on gene expression, we utilized single-cell eQTL datasets to determine whether the variant underlies cell-type-specific eQTLs^109^. Notably, our analysis revealed a pDC-specific eQTL for the rs2841277 variant in *PLD4* in SLE patient data (**Fig. 6h**).

**Fig 6.**
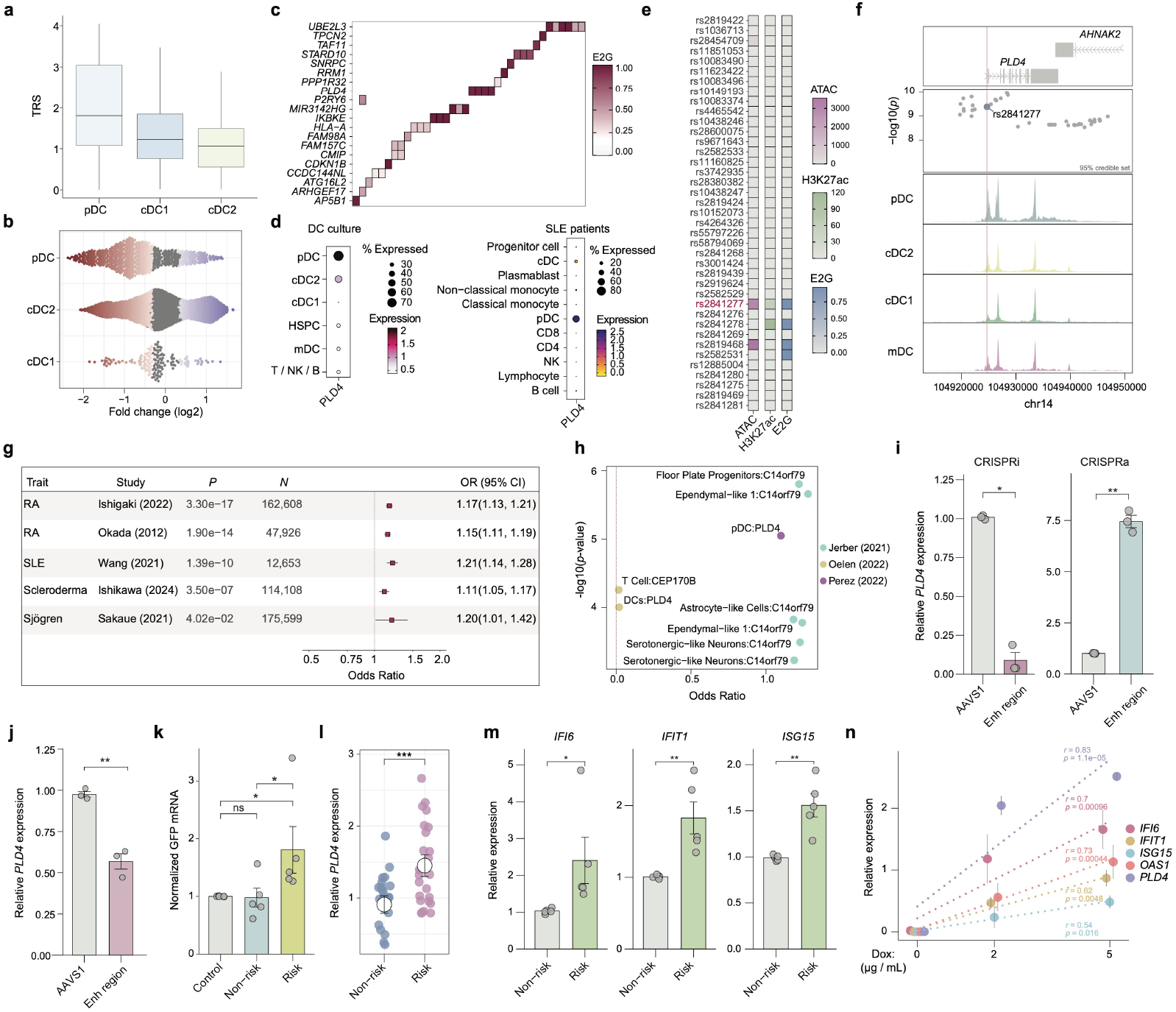
Functional assessment of SLE-associated genetic variants in pDCs. **(a)** Bar plot of the mean SLE TRS for each DC subset. **(b)**Beeswarm plot showing adjusted log2 fold-changes (log2FC) of cell population abundance between samples of SLE patients and healthy donors.**(c)** A heatmap of E2G scores for SLE associated variants-gene pairs overlapping a pDC enhancer or promoter. **(d)** Dot plot of *PLD4* expression across cell types in DC culture (left) or in SLE patients (right). Dot size indicates the percentage of cells expressing the gene, and color intensity represents the average gene expression. **(e)** SLE 95% credible set variants, with color indicating average ATAC-seq signal, H3K27ac enrichment, and E2G score. **(f)** Genome browser tracks showing cell-type-specific scATAC-seq profiles alongside GWAS summary statistics for SLE, with a zoomed-in view of the fine-mapped variants at the *PLD4* locus. **(g)** Forest plot showing the odds ratio and 95% confidence intervals for rs2841277 associations with different autoimmune diseases. **(h)** Scatter plot showing sc-eQTL analysis for rs2841277 and its linked gene in the indicated cell type and cohort. Statistical significance is shown as −log_10_(y-axis) relative to the effect size expressed as the Odds Ratio (x-axis).**(i)** Relative mRNA expression of *PLD4* after CRISPRi or CRISPRa targeting the *PLD4* regulatory element (‘Enh region’) compared to control(‘AAVS1’) in CAL-1 cells (*n* = 3 independent experiments). **(j)** Relative mRNA expression of *PLD4* in CAL-1 cells following CRISPR-Cas9-mediated microdeletion of the *PLD4* regulatory element (‘Enh region’) compared to an AAVS1 safe-harbor control (*n* = 3 independent experiments). **(k)** Allele-specific reporter assay for an rs2841277-centered fragment containing either the non-risk (C) or risk allele (T) in CAL-1 cells (*n* = 5 independent experiments). **(l)**, Relative mRNA expression of *PLD4* in HDR-edited isolated clones. **(m)** Relative mRNA expression levels of the indicated ISGs in PLD3^−/−^ CAL-1 HDR-edited clones stimulated with RNA9.2s for 4 h (*n* = 5 independent experiments). **(n)** PLD4^−/−^×PLD3^−/−^ CAL-1 cells were treated with the indicated doxycycline (Dox) concentrations and stimulated with RNA9.2s for 4 h. *PLD4* and ISG expression levels were quantified by RT-qPCR (*n* = 5 independent experiments). Data are shown as mean ± s.e.m. Statistical analysis was performed using a two-tailed, unpaired Student’s t-test (i, j, l, m). In (k), *P* values were determined by one-way ANOVA analysis followed by Tukey’s post hoc test for multiple group comparisons. In (n), two-sided P values and Pearson’s *r* were obtained from linear regression of relative mRNA expression versus doxycycline dose. \**P* < 0.05; \*\**P* < 0.01; \*\*\**P* < 0.001 or ns, not significant.

To functionally study the regulatory potential of this variant, we employed CRISPR interference and activation (CRISPRi/a) of the variant-harboring regulatory region (**Fig. S6f**) in the pDC-like cell line CAL-1. Analysis by qRT-PCR demonstrated that perturbation of this element led to significantly lower and higher *PLD4* expression levels by CRISPRi or CRISPRa, respectively (**Fig. 6i**). To further elucidate the putative impact of rs2841277, we used pairs of guide RNAs (gRNAs) to excise the SNP-harboring regulatory element (**Fig. S6f**). Consistent with our CRISPRi/a findings, targeted deletion of this region resulted in a reduction in *PLD4* levels (**Fig. 6j**). To evaluate whether enhancer activity was allelespecific, we compared the T risk allele to the C allele in a reporter assay. We observed increased reporter activity with the T risk allele (**Fig. 6k**). To enable precise genomic modifications, we employed Cas9-mediated homology-directed repair (HDR) to introduce single-base substitutions for the risk allele and the non-risk allele in CAL-1 single cell clones. Consistent with our reporter assay results, clonal lines carrying the risk allele exhibited higher *PLD4* expression (**Fig. 6l**). After establishing that this regulatory element controls *PLD4* expression, we investigated the functional requirement of *PLD4* for nucleic acid sensing. Given the reported functional redundancy between PLD3 and PLD4^104,110^, we knocked down the expression of *PLD3* in our HDR-edited cells. We found that clones carrying the risk allele exhibited a heightened response compared to clones carrying the nonrisk allele (**Fig. 6m**). To evaluate how *PLD4* levels independently modulate TLR7 responses, we generated *PLD4*x*PLD3* double-deficient CAL-1 cells (**Fig. S6g,h**). Consistent with previous reports, the loss of *PLD4* and *PLD3* resulted in a diminished response to stimulation with ssRNA, whereas the addition of exogenous *PLD4* rescued the functional deficits, restoring the induction of downstream interferon-stimulated genes (ISGs) (**Fig. S6i**). To further define this relationship, we used the TetON system to enable temporal control of *PLD4* expression. Under RNA stimulation, ISG induction increased in a *PLD4* dose-dependent manner, suggesting that elevated *PLD4* levels can lead to hyperactivation of TLR7-mediated signaling, shifting the early immune response toward a more rapid and robust activation profile (**Fig. 6n**). Finally, given that SCENIC+ identified TCF4 as the top candidate regulator of *PLD4* (**Fig. 5i** and **Fig S6j**), we performed siRNA-mediated knockdown of *TCF4*^111^ and observed a decrease in *PLD4* expression (**Fig. S6k**), supporting the model that *PLD4* is part of a TCF4-governed transcriptional program.

Collectively, our findings demonstrate that the *PLD4* risk variant rs2841277 drives pDC-specific disease predisposition to developing SLE and other autoimmune disorders, identifying a cell-type-specific regulatory mechanism that increases TLR7-mediated responses and illustrating how our single-cell multi-omic atlas can enable mapping of key disease mechanisms in complex immune disorders.

## Discussion

Here, we establish a comprehensive map of human DC differentiation from HSPCs to all DC subtypes at single-cell resolution, enabling the reconstruction of DC gene regulatory networks and filling fundamental gaps in our understanding of human DC development. We provide an integrated analysis across multiple tissues to dissect the conservation in human DC transcriptomes among various subtypes.

Our multi-omic atlas provides insights into cell-type-specific GRNs. These data support and extend findings from mouse models to human DC development. In line with mouse studies where Zeb2 ablation increases splenic cDC1 populations^33^, we demonstrate that ZEB2 regulates cDC1 specification through its antagonistic relationship with ID2. However, while Zeb2 is required by mouse pDCs^34^, human pDCs were unaffected in our model. Within committed DC progenitors, the branching of pDC and cDC1 is determined by the balance between ID2 and TCF4. We show that genetic perturbation of *TCF4* results in reduced pDC numbers and a moderate increase in cDC1s. Consistent with our findings in human cells, *Tcf4* deletion in mice impaired pDC development. While mouse cDC lineages remain largely unaffected, *Tcf4* deletion led to the expansion of the CD8+ cDC phenotype likely through Id2 induction^45,112^. Our mtDNA-based clonal inference further supports the role of this TCF4/ID2 antagonism model in pDC/cDC1 divergence, akin to what has been previously described in mice^48^.

In many GWASs, the relevant cell types or states, as well as underlying mechanisms, through which genes confer disease risk remain to be identified^113^. Interpretation of regulatory landscapes from large single-cell epigenomic atlases must account for the disproportionate statistical power of the most abundant cell types, which can bias interpretation in GWAS functional follow-up^114^. For rare cell populations, the limited number of cells per donor often necessitates ‘pseudobulk’ aggregation that may not meet the minimum signal-to-noise thresholds required for accurate peak identification within a population. Consequently, downstream enrichment analyses may be inherently biased, underrepresenting the regulatory logic of rare, but biologically pivotal, cell states^115^. We provide a key example of this through our studies of rare DC subtypes, as many of the observed complex disease associations might be missed in conventional analyses without producing a resource with deep single-cell multi-omic coverage across the diversity of human DC subtypes. Indeed, we connect a variety of complex immune diseases with specific DC subtypes, enabling a range of future studies seeking to dissect underlying mechanisms.

IFN dysregulation by pDCs plays a primary role in the pathogenesis of SLE by regulating the differentiation of autoantibody-producing B cells^116,117^. However, the triggers of these uncontrolled type I IFN responses, particularly the inherited genetic predisposition leading to pDC dysregulation, and their relationships with SLE development remain largely undefined^118^. Transitioning from association to causality requires functional dissection of GWAS-implicated variants. Utilizing functional genomics tools such as SCAVENGE and scE2G mapping, we demonstrate that common variants associated with SLE are significantly enriched within pDC-specific regulatory elements. Specifically, we identify the *PLD4* locus as a prime example of how developmental and cell-type-specific epigenomic data can reveal previously hidden drivers of disease. We provide a deeper mechanistic understanding of the diverse roles of pDCs in immunity by linking SLE-associated genetic variants to the pathways that drive autoimmunity, thus expanding our understanding of the functional consequences of this variation. Targeting such pathogenic variants that modulate TLR signaling or other inflammatory pathways in pDCs could enable new therapeutic approaches for SLE treatment and prevention.

Our data enable the systematic mapping of genetic variation underlying immune-related and other diseases that extend beyond genetic associations, particularly those that occur in rare and hard to study cell types, offering insights into the mechanisms of human diseases that can be exploited for new therapeutic avenues.

## Acknowledgements

We thank members of the Sankaran laboratory for valuable feedback. This work was supported in part by the Myasthenia Gravis Foundation of America High-Impact Pilot Project Award (to V.G.S), the National Eczema Association Catalyst Research Grant no. NEA23-CRG185 (to O.C.), the Howard Hughes Medical Institute (to V.G.S.), Alex’s Lemonade Stand Foundation (to V.G.S.), Blood Cancer United (to V.G.S.), the Edward P. Evans Foundation (to V.G.S.), the Gates Foundation (to V.G.S.), and the National Institutes of Health Grants (R01DK103794, R01CA265726, R01CA292941, R33CA278393, R01HL146500). A.-L.N is supported by an EMBO Postdoctoral fellowship (ALTF 209-2024). V.G.S. is an investigator of the Howard Hughes Medical Institute.

## Author contributions

O.C. and V.G.S. conceived and designed the study. O.C led the computational analyses and performed the experiments. C.W, T.Y, A.-L.N, C.G provided valuable input on experimental design. L.B.N and E.K assisted with the data analyses. O.C and V.G.S wrote the manuscript with input from all authors. V.G.S. provided overall project oversight and acquired funding.

## Materials and Methods

### Cell line and primary cell culture

Human umbilical cord blood was obtained from Brigham and Women’s Hospital as discarded deidentified samples. CD34^+^ HSPCs were enriched using the MicroBead Kit UltraPure CD34^+^ positive selection (Miltenyi Biotec) according to the manufacturer’s instructions. For cord blood culture, MS5 cells (DSMZ) were maintained in complete α-MEM medium (Invitrogen) with 10% fetal calf serum (FCS) and penicillin/streptomycin (Invitrogen). After 3 h of 10 µg/mL mitomycin C (Sigma-Aldrich) treatment and washing, 3 × 10^4^ MS5/mL were seeded per well in 96-well plates 24 h before culturing hematopoietic cells. Bulk CD34^+^ HSPCs were seeded in a medium containing 100 ng/mL FLT3L (PeproTech), 20 ng/mL SCF (PeproTech), and 10 ng/mL GM-CSF (PeproTech). Cells were harvested between days 14-17 for flow cytometry analysis.

CAL-1 cells (DSMZ) were cultured in RPMI 1640 medium supplemented with 10% heat-inactivated FCS and 100 U/mL penicillin/streptomycin.

### Single-cell multiome and mtDNA-enriched library generation

Lineage^−^CD45^+^HLA-DR^+^ cells were sorted into collection tubes with 2% FCS. Approximately 15,000 nuclei were loaded into a Chromium Controller to produce single-cell gel beads, following the single-cell Regulatory Multiomics (transcriptomics and chromatin accessibility) with Deep Mitochondrial Mutation Profiling (ReDeeM) protocol^2323^. Briefly, cells were fixed, permeabilized (0.1% formaldehyde, 0.04% BSA, 0.2 U/µL RNase inhibitor in DPBS) and tagmented (ATAC buffer B; PN-2000193, ATAC Enzyme B; PN-2000265 and glyco-diosgenin; Avanti Polar Lipids, 850525P), followed by GEM generation, barcoding, post-GEM cleanup, pre-amplification PCR, cDNA amplification, and library construction using the 10x Genomics Chromium Next GEM Single Cell Multiome ATAC + Gene Expression protocol (CG000338, Rev C). mtDNA sequences were enriched from ATAC sequencing libraries by hybridization capture using a custom staggered probe strategy. For each sample, a total of 2,000 ng of ATAC library was equally divided across four independent hybridization reactions, each utilizing a distinct custom probe panel. Hybridization capture and subsequent washes were performed using the xGen Hybridization and Wash Kit (IDT) according to the manufacturer’s protocol. Following capture, post-capture PCR amplification was performed and monitored by SYBR Green. The four amplified reactions corresponding to each sample were subsequently pooled and purified to yield the final enriched mtDNA library.

### Single-cell multiome data processing and analysis

To generate count matrices for both RNA and ATAC modalities, raw sequencing data were processed using the Cell Ranger ARC pipeline (v.2.0.2) with alignment to the human reference genome (GRCh38). Cell Ranger ARC output files were processed using the Seurat and Signac R packages (versions 5.4 and 1.16). QC filtering was applied using the following thresholds: nCount_ATAC > 120,000, TSS.enrichment > 1, nucleosome_signal < 2 and nCount_RNA > 1,000. RNA data were normalized and scaled using the SCTransform function, followed by principal component analysis and uniform manifold approximation and projection (UMAP) for dimensionality reduction. ATAC data were normalized using term frequency-inverse document frequency (TF-IDF) normalization, followed by singular value decomposition and the UMAP function in Signac. To integrate the RNA and ATAC modalities, we performed weighted nearest neighbor (WNN) analysis using Seurat’s FindMultiModalNeighbors function to construct a WNN graph. The datasets from three different donors were then integrated using SelectIntegrationFeatures, PrepSCTIntegration, FindIntegrationAnchors, and IntegrateData, with default parameters. Cell types were annotated based on the expression of canonical marker genes identified using a Wilcoxon rank sum test as implemented in the presto package. Motif activities were calculated using Signac’s RunChromVAR function and the JASPAR2024 database.

### Single-cell mtDNA clonal inference

For each donor, a single-cell lineage tree was reconstructed from mtDNA variant allele counts using MitoDrift with 10 Markov chain Monte Carlo (MCMC) chains and a confidence threshold (τ = 0.2). Cell clusters were grouped into major DC subsets (cDC1, cDC2 and pDC) to ensure sufficient cell counts for each population. To quantify clonal sharing, a binary presence/absence matrix of clone-to-subtype pairings was generated. Lineage coupling between DC subsets was evaluated using the Szymkiewicz–Simpson overlap coefficient. This metric is defined as the size of the clonal intersection divided by the size of the smaller of the two compared sets, which controls for differences in cell numbers.

### Cell stimulations

HSPC-derived DCs were stimulated with R848 (2.5 µg/mL, InvivoGen) or lipopolysaccharide (LPS) (10 ng/mL, L2654, Sigma-Aldrich). After 16 h of incubation, DCs were stained antibodies against CD83 (HB15e, FITC, BioLegend) and CD86 (W17233E, Brilliant Violet [BV] 421, BioLegend) and analyzed by flow cytometry.

CAL-1 cells were primed with hIFN-γ (10 ng/mL, PeproTech) for 30 min prior to and during stimulation (4 h). For transfection with RNA9.2s (rArGrCrUrUrArArCrCrUrGrUrCrCrUrUrCrArA, 0.65 µg/well), RNA and poly-L-arginine (P7762, Sigma-Aldrich) were incubated separately in a 1:1 ratio for 5 min in pre-warmed Opti-MEM (25 µL/well). Subsequently, the two reagents were combined, incubated for an additional 20 min, and added to the cells

### Antibodies and reagents

For purification of differentiated DCs, cells were stained with LIVE/DEAD Fixable Blue stain (Life Technologies), CD45 (HI30, Alexa Fluor 700 [AF700]; BioLegend), CD15 (HI98, APC-Cy7; BioLegend), CD56 (HCD56, APC-Cy7; BioLegend), CD19 (HIB19, APC-Cy7; BioLegend), CD3 (OKT3, APC-Cy7; BioLegend), CD20 (2H7, APC-Cy7; BioLegend), CD14 (TuK4,Qdot-655; Invitrogen), CLEC9A (9A11, PE; eBioscience), CD1c (L161, PE-Cy7; BioLegend), CD303 (201A, FITC; BioLegend), CD304 (12C2, FITC; BioLegend), CD123 (6H6, BV510; BioLegend), CD141 (AD5-14H12, APC;Miltenyi Biotec) and HLA-DR (L243, AF700, BioLegend) for 30 min on ice. For flow cytometry analysis, cDC1s (Lin^−^HLA-DR^+^CLEC9A^+^CD141^+^), cDC2s (Lin^−^CD14^−^HLA-DR^+^CD1c^+^) and pDCs (Lin^−^CD1c^−^CD14^−^HLA-DR^+^CD303/4^+^CD123^+^) were analyzed or sorted with a FACSAria IIIu in-strument (BD Biosciences). Data were collected with the FACSAria IIIu instrument and analyzed by FlowJo (v10.2).

### Morphological analysis

Purified DCs were analyzed by Giemsa staining of cytospin preparations. As few as 5 × 10^4^ cells were cytospun for 5 min at 800 rpm on a glass slide and then stained with the Hemacolor stain kit (Harleco) as recommended by the manufacturer. Slides were then imaged on an Axioplan 2 microscope (Carl Zeiss) at 100× magnification.

### Ectopic expression of *PLD4*

For the expression of PLD4 (LeGO-iG2-PLD4), the *PLD4* cDNA fragment was subcloned into the LeGO iG2 vector (Addgene #27341). CAL-1 cells were transfected at a density of 1 × 10^6^ cells/well with *PLD4* plasmid or empty plasmid using the 4D-Nucleofector System with the DN100 (cell line SF, Lonza) program.

For the construction of inducible *PLD4*, the pLVX-TetOne backbone was amplified from TetOne-Puro-GFP (Addgene #171123) and *PLD4* cloned into the AgeI and EcoRI restriction sites. *PLD4* expression was induced by adding 2 or 5 µg/mL doxycycline (D9891, Sigma-Aldrich) for 48 h before stimulation.

### Trait enrichment analysis

GWAS datasets for inflammatory, autoimmune, and oncologic diseases were selected from FinnGen (R9)^68^ and CAUSALdb^48^ (Suppl. Table 5). Subsequently, we retrieved the 95% credible sets of fine-mapping results of each GWAS with the posterior probabilities generated by FINEMAP^119^. We identified trait-relevant cell types using SCAVENGE in conjunction with our scATAC-seq data and GWAS loci. We used the fine-mapping results and scATAC-seq from all clusters as inputs for SCAVENGE, which assigned a trait relevance score (TRS) to each cell type. Significant trait-enriched cells (*P* < 0.05) were identified by comparing their network propagation scores against an empirical null distribution, which was generated from 1,000 permutations using degree-matched random seed cells.

For g-chromVAR^120^ analysis, we summed the accessibility profiles of major DC subsets to create a pseudobulk accessibility matrix. To ensure stability, we ran g-chromVAR 50 times using default parameters and averaged the results.

### Gene editing of cells

For CRISPR interference (CRISPRi) and activation (CRISPRa), the dCas9-KRAB coding region (Addgene #220838) and the dCas9-VP64 coding region were individually subcloned into the PEmax-mRNA (IVT) backbone (Addgene #204472). *In vitro* transcription was then performed to generate purified mRNA encoding dCas9-KRAB or dCas9-VP64. CAL-1 cells were washed twice in DPBS and resuspended in 20 µL of Lonza SF buffer with supplement.

For each nucleofection, cells were mixed with 2 µg dCas9-KRAB or dCas9-VP64 mRNA and a total of 2 µL sgRNA targeting either the AAVS1 safe-harbor locus or the rs2841277-harboring regulatory element with independent pairs of sgRNAs. Cells were electroporated in 20 µL Nucleocuvette strips using the Lonza 4D-Nucleofector System, with the DN100 program. Immediately after electroporation, 75 µL of prewarmed medium was added to the electroporation cuvette, which was placed in an incubator at 37 °C for 5 mins. Cells were then plated at a density of 2.5×10^5^ cells/mL in adequate complete medium. 72 h after nucleofection, total RNA was extracted, cDNA was generated, and real-time quantitative PCR (RT-qPCR) was performed to detect *PLD4* expression.

To form the ribonucleoprotein (RNP) complexes, 100 pmol of Alt-R V3 Cas9 protein (IDT) was incubated with 100 pmol of chemically modified sgRNA (Synthego) for 20 min at room temperature. Primary human CD34^+^ HSPCs, cultured for 36–48 h, were harvested and resuspended in 20 µL of P3 Primary Cell Nucleofector Solution (Lonza) supplemented with 1 µL Electroporation Enhancer (IDT). The cell suspension was combined with the preformed RNP complexes and transferred to a nucleocuvette strip. Electroporation was performed using the 4D-Nucleofector System (Lonza) under program DZ-100.

For targeted genomic editing via homology-directed repair (HDR), RNP complexes were prepared by incubating 100 pmol Alt-R V3 Cas9 protein with 100 pmol cutting sgRNA (IDT) for 15 min at room temperature. To serve as a repair template, a 200-nucleotide single-stranded oligodeoxynucleotide (ssODN) was synthesized (IDT). The donor oligo was designed with 90-nt homology arms flanking the variant target site and incorporated phosphorothioate linkages at both termini to increase stability. CAL-1 cells were resuspended in 20 µL of SF nucleofection solution supplemented with 1 µL of Electroporation Enhancer (IDT). The cell suspension was combined with the pre-formed RNP complex, followed by the addition of 0.5 µL of 100 µM donor oligo (1:50 dilution). Electroporation was performed using the 4D-Nucleofector System with program DN-100. Immediately following nucleofection, cells were recovered in culture media supplemented with AltR HDR Enhancer V2 (IDT). After 24 h, cells were washed with PBS and transferred to standard culture media for further expansion and downstream analysis of editing efficiency.

### Reporter assay

A 300-bp fragment centered on the variant was amplified directly from CAL-1 cell genomic DNA and subjected to In-fusion assembly into SbfI/AgeI-digested pLS-mP (Addgene #81225). As a negative control, part of the *ACTB* genomic locus was cloned. Oligonucleotides used for the PCR amplification are shown in Suppl. Table 6. The production of lentiviral particles containing reporter sequences was performed as described above. To analyze reporter expression, genomic DNA and total RNA were extracted from the transduced cells, followed by reverse transcription of RNA into cDNA and analysis by RT-qPCR. eGFP mRNA and DNA levels were normalized using *ACTB* as a reference gene, and enhancer activity was calculated as (normalized eGFP mRNA level)/(normalized eGFP DNA level).

### Cell types similarities

Public single-cell RNA-seq datasets (Suppl. Table 1) were analyzed in Seurat. Original cell-type annotations were retained, and datasets were subset to DC lineages and normalized using SCTransform, followed by PCA and UMAP. Cell-type transfer from the *in vitro* DC dataset was performed with FindTransferAnchors and TransferData functions, and predicted labels were added to each query object for comparison. Cross-dataset concordance was evaluated by comparing transferred labels to original annotations and summarizing the distribution of predicted identities within each annotated cell type.

Integration of scRNA-seq datasets and generation of a cell-to-cell-type similarity matrix were performed using CellHint^121^. Cell types were annotated using CellTypist^28^.

### GRN inference and trajectory analysis

Gene regulatory networks were inferred using the Pando^30^ and SCENIC+^37^ frameworks. Initially, potential regulatory links were identified using the infer_grn function from Pando, with default parameters. To refine the network, GRN edges were filtered into functional modules using the find_modules function. To identify cell-type-specific GRNs, the network was further constrained by incorporating differentially accessible regions (DARs). Only regulatory interactions involving peaks identified as differentially accessible between specific cell types were retained for the final cell-type-specific models.

Differentiation states among DC subpopulations were inferred using the CytoTRACE2 R package (v1.1)^29^ with default parameters. To reconstruct differentiation dynamics, we used spliced and unspliced transcripts (velocyto v0.17.17)^32^ to recover the latent time using the scVelo package^31^ (v0.3).

### Reverse transcription quantitative PCR (RT-qPCR)

Total RNA was extracted with the Total RNA Purification Micro Kit (Norgen, 35300) according to the manufacturer’s protocol. Reverse transcription (500-1,000 ng total RNA) was conducted using the PrimeScript RT reagent Kit (#RR037B; TAKARA). Reverse transcription quantitative polymerase chain reaction (RT-qPCR) was conducted using SYBR Green (A25743, Thermo Fisher Scientific, USA) for specific primers to *ACTB, PLD4, IFI6, TCF4, ISG15, OAS1* and *IFIT1* (Suppl. Table 6).

### Linking variants to target genes

To map associated variants to effector genes we intersected GWAS variants with the single-cell enhancer-to-gene (scE2G)^72^ framework. Variants were further filtered based on differential expression and relevance of their target genes within the DC subset.

### Single-cell differential abundance

We used the MiloR package (v1.2.0) to perform differential abundance analysis in cells within the inferred neighborhoods, between SLE patients and healthy controls. We first used the buildGraph function to construct a KNN graph (k=30, d=15-30). Next, we used the make_Nhoods function to assign cells to neighborhoods based on their connectivity over the KNN graph. The neighborhoods were projected onto the UMAP embedding for visualization. To test for differential abundance, Milo fit a Negative Binomial GLM to the counts of each neighborhood, accounting for different numbers of cells across samples using the trimmed mean of M-values (TMM) normalization. The spatial FDR and log_2_FC of the number of cells between two conditions in each neighborhood were used for visualization. The beeswarm plot of the distribution of log_2_FC across neighborhoods was used to present the differential abundance of cell populations.

We also used the sccomp framework for differential composition and variability analysis. Predictive posterior intervals were checked after fitting to assess the model’s descriptive adequacy to the data. sccomp identified outlier observations (cell-type abundance) and dropped them from the hypothesis testing.

### sc-eQTL data

To investigate the cell-type-specific effects of the rs2841277 variant across immune and non-immune lineages, we utilized data from scQTLbase^109^, a comprehensive repository of human single-cell eQTLs.

### Statistics and reproducibility

For statistical evaluation, an unpaired Student’s *t*-test or one-way ANOVA with Tukey’s post *hoc test* for multiple comparisons was applied for calculating statistical probabilities in this study. For all panels, at least three independent experiments were performed. Data were analyzed using R. *P* values smaller than 0.05 were regarded as statistically significant. * *P* < 0.05, *P* < 0.01, *** *P* < 0.001, **** *P* < 0.0001; ns, not significant.

## Data availability

All data and code will be released at the time of publication.

---

**Supplemental Fig. 1:**
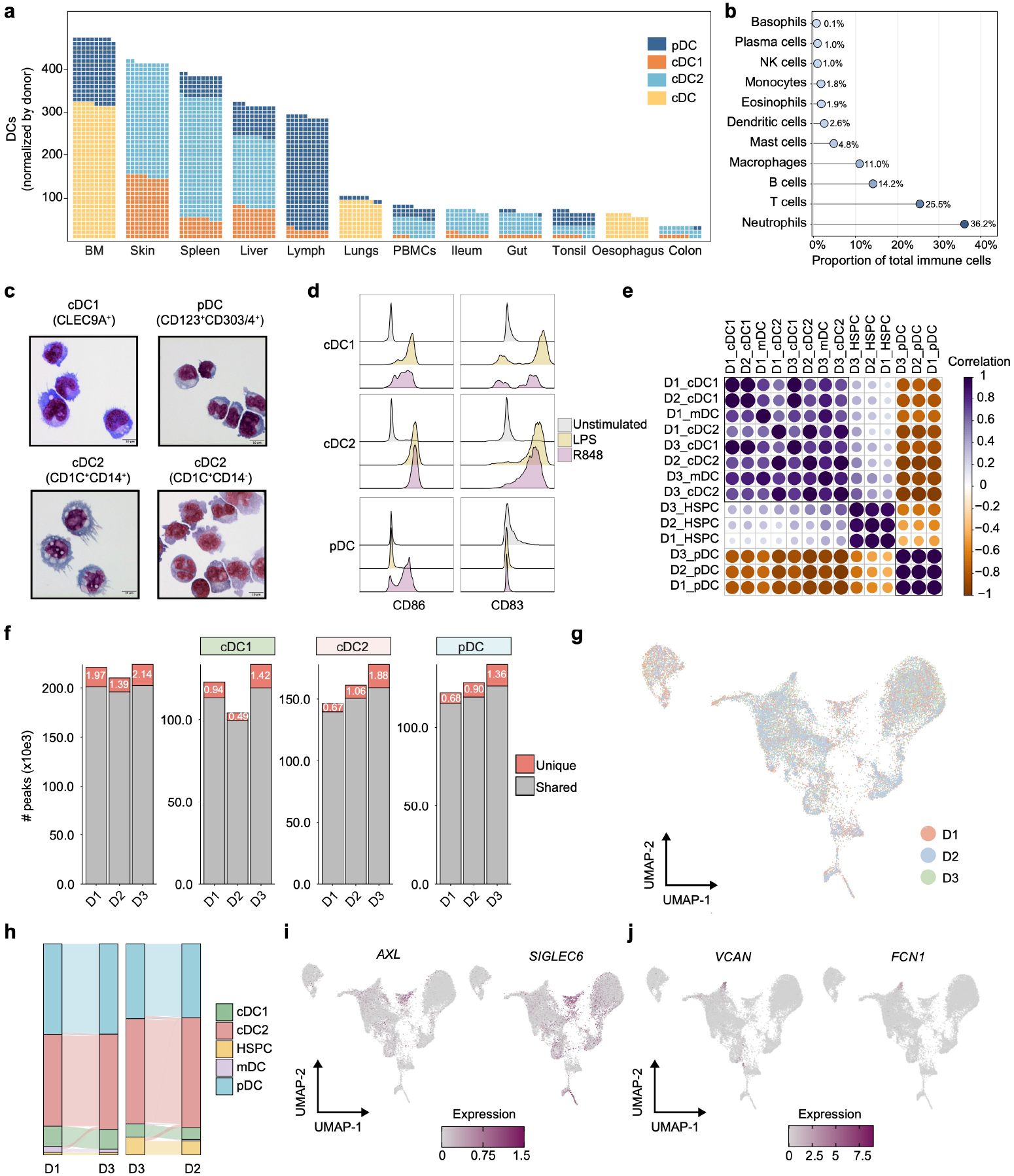
Concordant among donors. **(a)** Average DC numbers per donor obtained from scRNA-seq datasets (Suppl. Table 1). **(b)** Lollipop plot showing the proportion of total immune cells represented by each immune cell type, based on estimated total cell numbers (adapted from Sender *et al*., 2023)^13^. **(c)** May-Grünwald-Giemsa staining of sorted DC subsets. Scale bars, 10 μm. **(d)** Activation marker expression of CD83 and CD86 on the indicated DC subtypes 16 h after LPS or R848. **(e)** Correlation of the mean values of motif activity calculated by chromVAR for each cell type across donors. **(f)** Comparison of scATAC-seq peaks that were accessible in all donors and overlapping across major DC subtypes. **(g)** Integrated UMAP of the three donors. **(h)** Sankey diagram showing the transcriptomic mapping of major cell types across three independent donors. **(i)** Average expression of tDC markers projected on the UMAP plot. **(j)** Average expression of DC3 markers projected on the UMAP plot. Abbreviations: BM, bone marrow.

**Supplemental Fig. 2:**
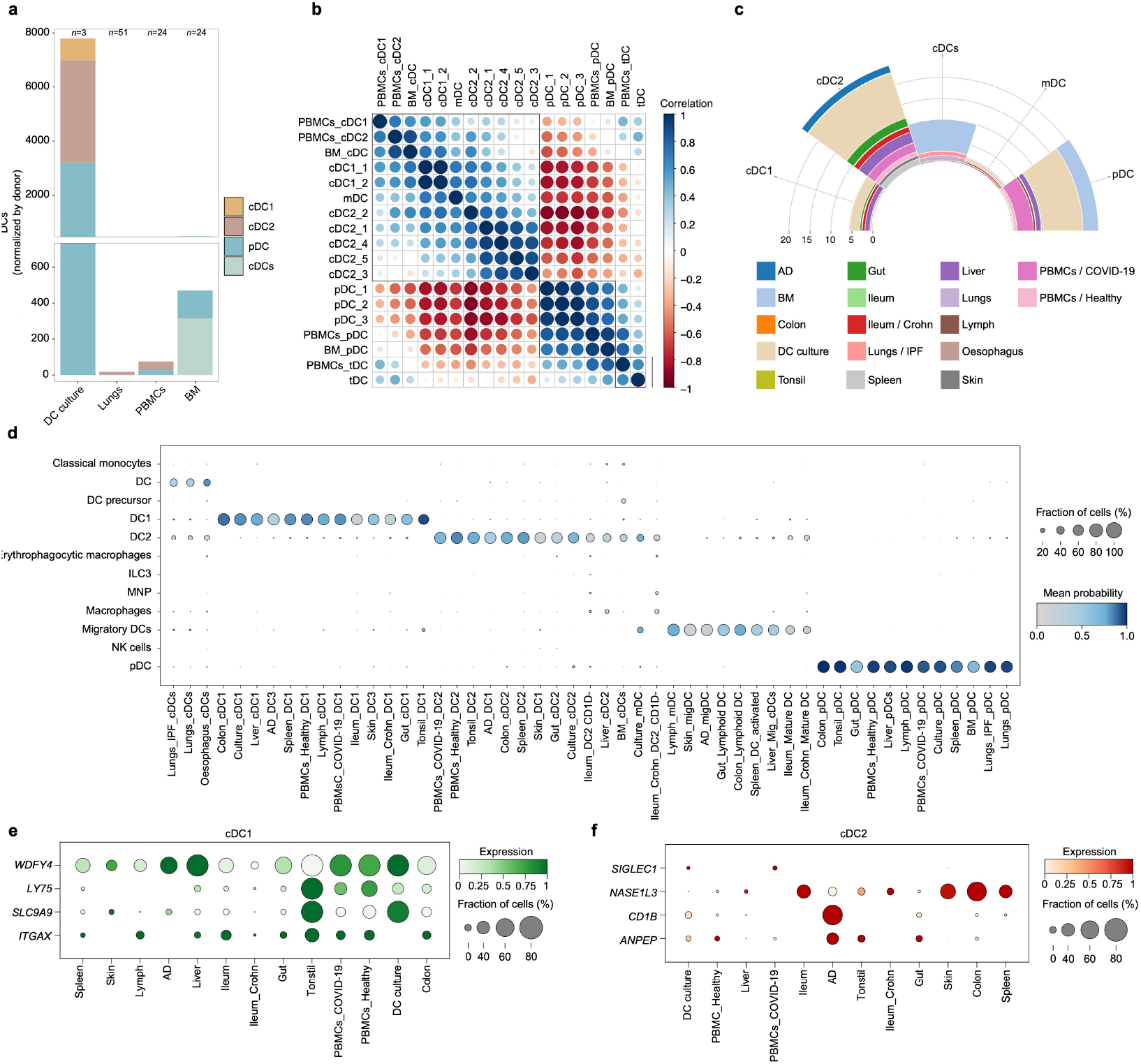
Comprehensive characterization of DC subsets across tissues. **(a)** Average DC numbers per donor obtained from the indicated datasets (Suppl. Table 1). **(b)** Correlation of mean chromVAR motif activity per cell type from a single donor across datasets (DC culture, PBMCs, and BM)^23,122^. **(c)** Cell counts and distribution of DC subsets across each integrated dataset. **(d)** DC subpopulations annotated using the CellTypist immune model. The color scale represents the mean probability score for each annotation. **(e,f)** Dot plots showing the expression of representative tissue-specific genes across the indicated DC subtypes and tissues of origin. Dot size indicates the percentage of cells expressing the indicated gene, and color intensity represents the average gene expression. Abbreviations: BM, bone marrow; AD, atopic dermatitis; IPF, idiopathic pulmonary fibrosis.

**Supplemental Fig. 3:**
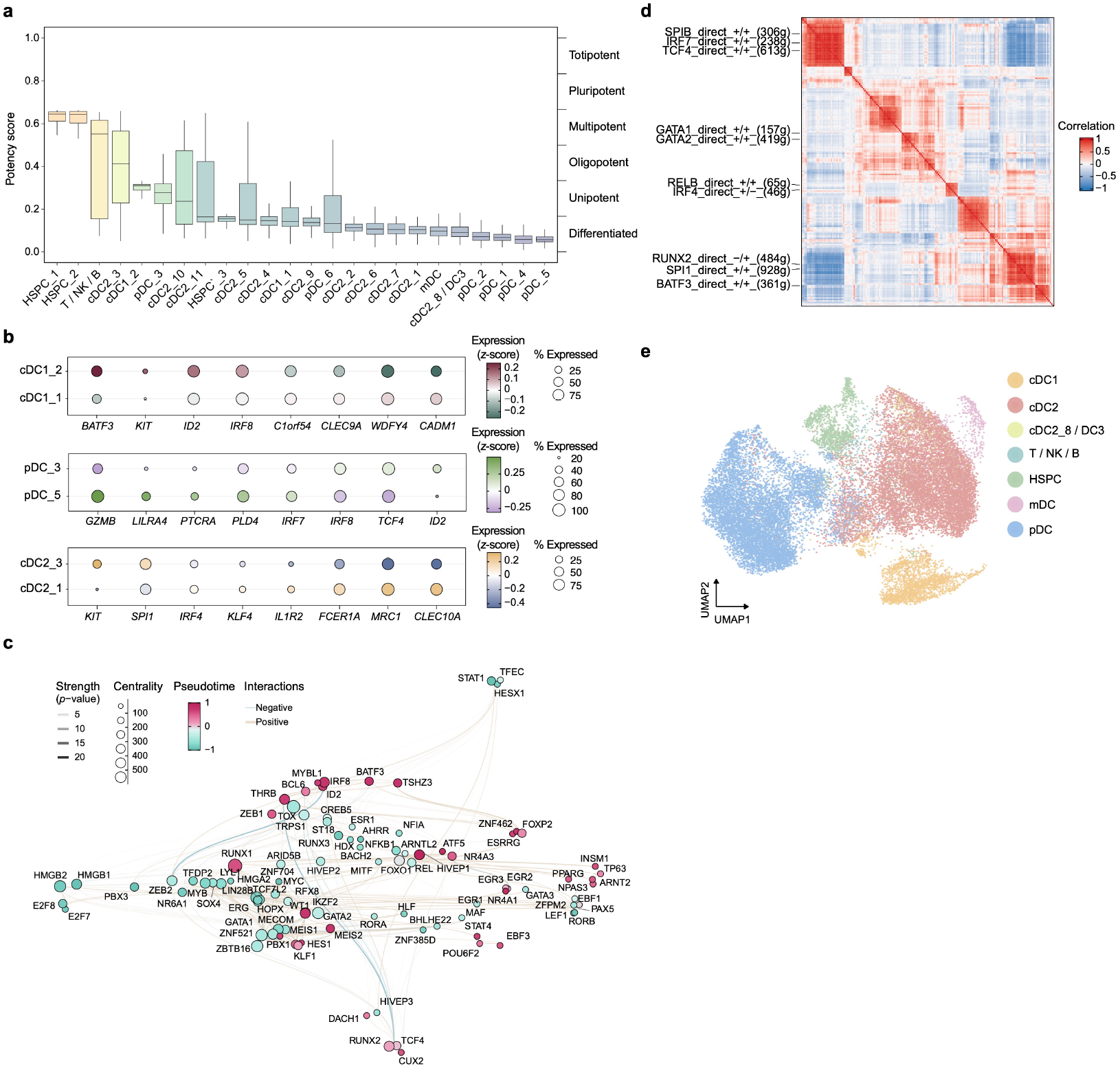
Developmental potency of human DC differentiation. **(a)** Box plots of the potency scores of each cell cluster calculated by CytoTRACE2 and ordered by potency. **(b)** Dot plot showing marker gene expression across early and late subsets ordered by progeny rank in **a**. Dot size indicates the percentage of cells expressing each gene. Dot color represents the average *z*-scored expression within each group. **(c)** GRN UMAP embedding of the inferred TF modules based on co-expression and interaction strength between TFs. Node size represents the number of connections for each TF. Color represents gene expression correlation with pseudotime (−1, early; 1, late). **(d)** Correlation between SCENIC+ eRegulons. **(e)** UMAP embedding based on target gene and target region enrichment scores of SCENIC+ eRegulons.

**Supplemental Fig. 4:**
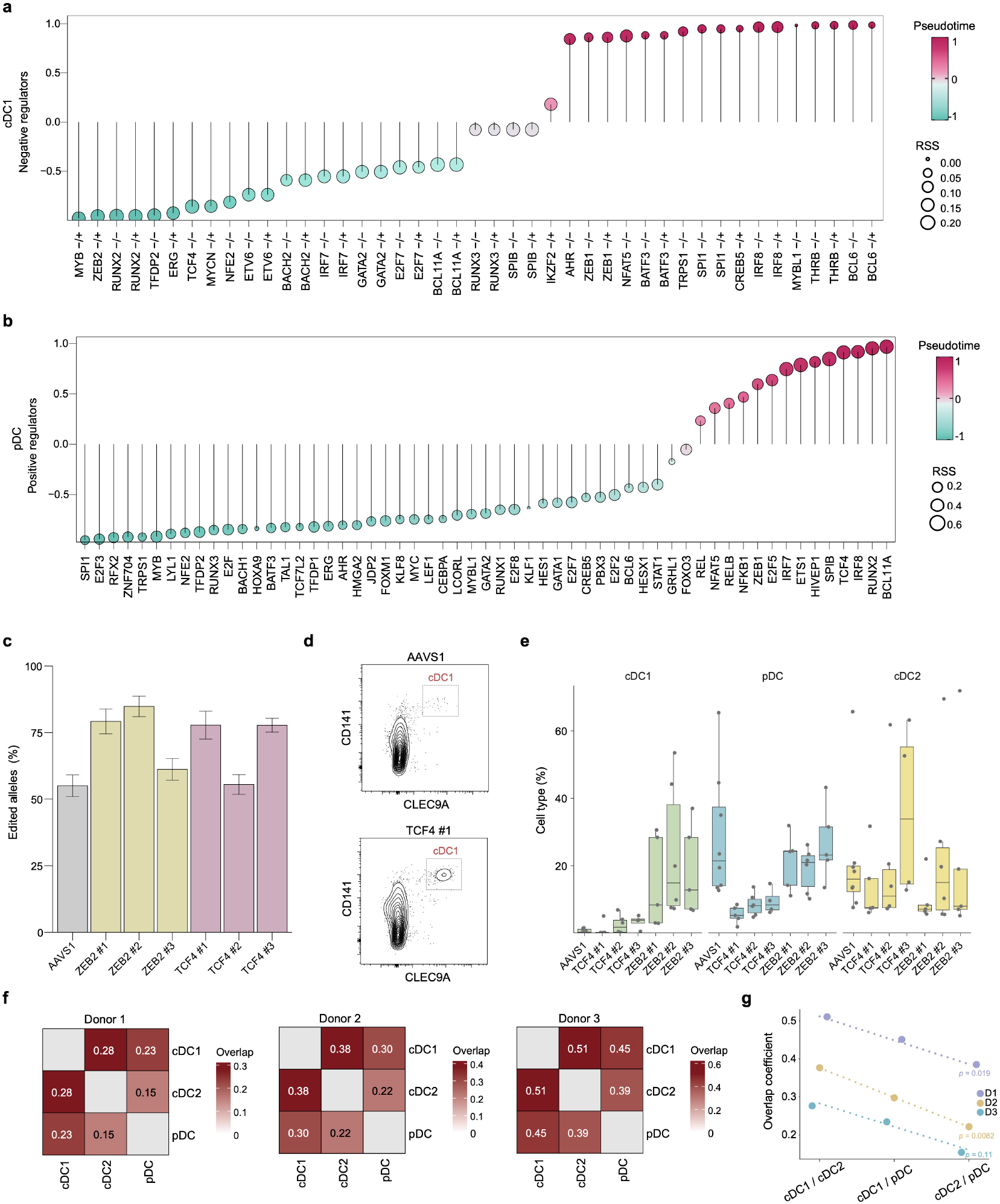
Regulatory landscape and genetic perturbation of human DC subsets. **(a,b)** Lollipop plots showing TF expression correlation with pseudotime (−1, early; 1, late) on a color scale and regulon specificity for cDC1s (**a**) and pDCs (**b**). **(c)** Editing efficiency, measured as the indel frequency of a given sgRNA. **(d)** Representative flow cytometry analysis of the cDC1s following CRISPR-Cas9-mediated knockout of *TCF4* or *AAVS1* (control). **(e)** Box plots showing the proportions of the indicated DC subsets following CRISPR/Cas9-mediated knockout of *TCF4, ZEB2*, or the control locus *AAVS1* (*n* = 8 independent donors). **(f)** Heatmap showing the MitoDrift-inferred clonal overlap among cDC1, cDC2, and pDC subsets. Color represents the overlap coefficient (Szymkiewicz–Simpson) for a given pair. **(g)** Trend analysis across lineage pairs (cDC1/cDC2, cDC1/pDC, and cDC2/pDC) for three independent donors. Individual data points represent the overlap coefficient (Szymkiewicz–Simpson) for a given pair. Dashed lines indicate donor-specific linear regression trends, with corresponding Pearson correlation (*r*).

**Supplemental Fig. 5:**
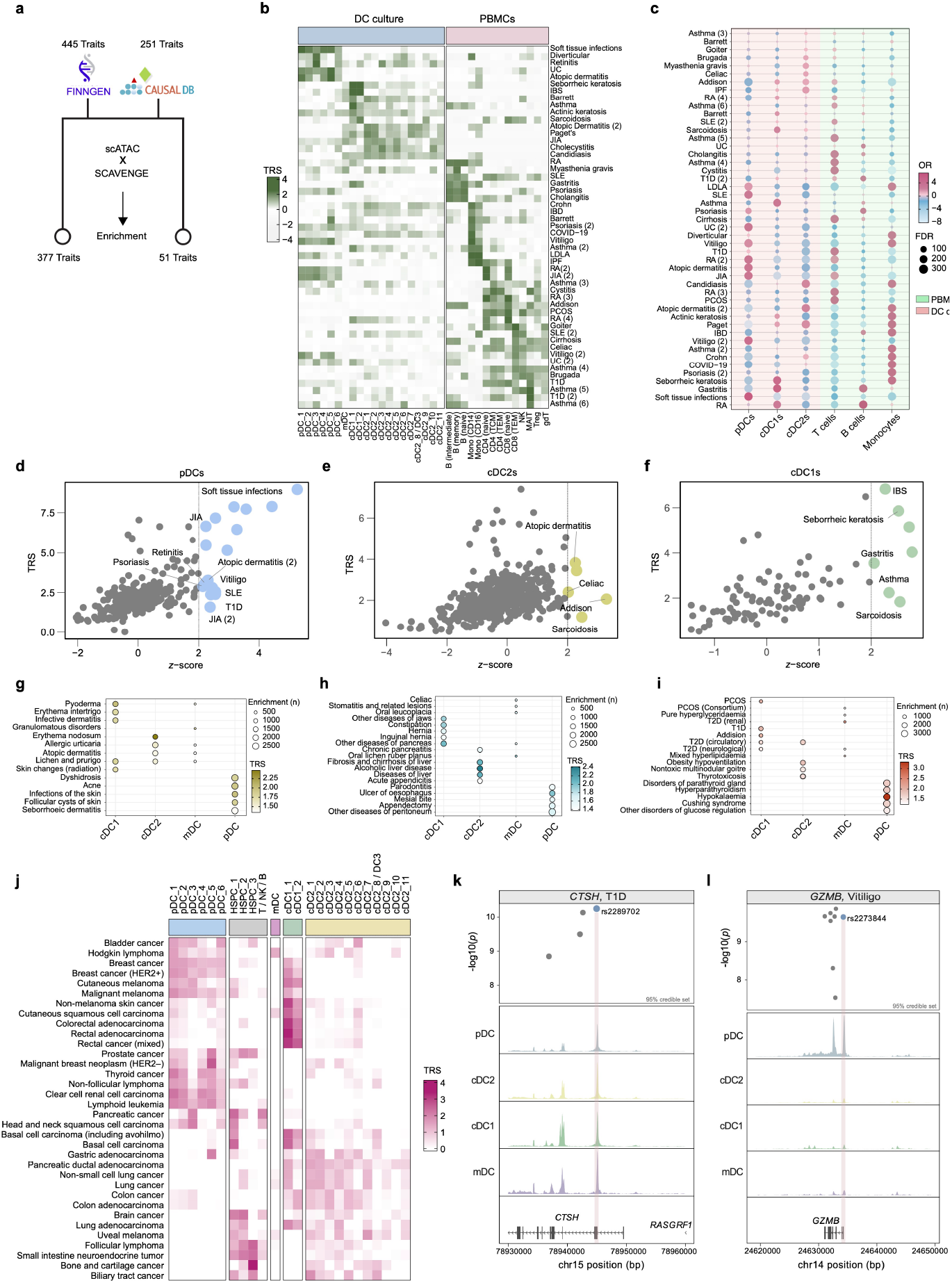
Cell-type-specific disease enrichment analysis. **(a)** The plot shows the number of autoimmune and inflammatory traits curated from FinnGen (R9) and CAUSALdb that were utilized for TRS calculation via SCAVENGE. (**b)** Heatmap showing TRSs (*z*-scores) for each cluster in integrated scATAC-seq profiles from DC culture and PBMCs^120^. **(c)** Bubble diagram for TRS enrichment analysis. Bubble size indicates the significance of enrichment (*FDR* < 0.05, two-sided hypergeometric test), scaled by −log_10_(*FDR*). Bubble color denotes the effect size log_2_(OR). Background shading indicates the data source: PBMCs (green) or in *vitro*-derived DC culture (red). **(d-f)** Enrichment of the indicated subtypes in complex diseases using SCAVENGE (y-axis) and gChromVAR (x-axis). Significant cell-specific enrichment is defined as *z*-score > 2 over the pseudobulk accessibility matrix. (**g-i**) FinnGen cohort: Heatmap of TRSs (*z*-scores) and the number of enriched cells for each cell type in diseases of the(**g**) skin, (**h**) digestive system, and (**i**) endocrine, nutritional, and metabolic systems. **(j)** Heatmap showing the TRSs (*z*-scores) of the indicated cancers for each cell type. **(k, l)** Normalized scATAC-seq signal coverage tracks showing cell-type-specific chromatin accessibility alongside GWAS summary statistics for the indicated traits, with a zoomed-in view of fine-mapped variants at the loci. Abbreviations: IBS, irritable bowel syndrome; T1D, type 1 diabetes; PCOS, polycystic ovary syndrome; IBD, inflammatory bowel disease; IPF, idiopathic pulmonary fibrosis; JIA, juvenile idiopathic arthritis; UC, ulcerative colitis; LADA, latent autoimmune diabetes in adults.

**Supplemental Fig. 6:**
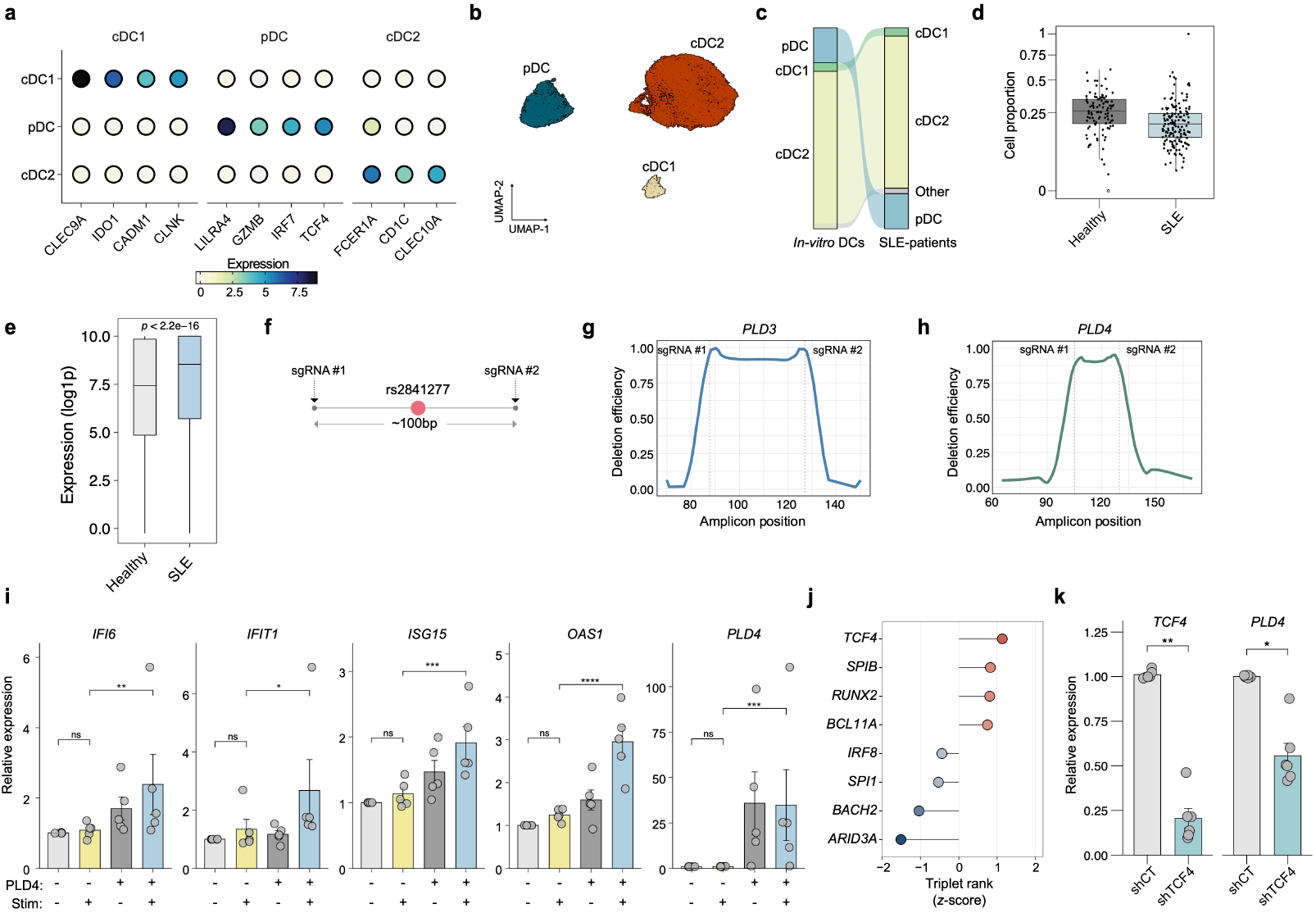
Characterization of fine-mapped SLE variants. **(a)** Average expression of DC markers derived from patients with SLE (PBMCs). (**b**) UMAP of DC populations derived from patients with SLE (PBMCs). **(c)** Sankey diagram showing the transcriptomic mapping of circulating DC populations from SLE patients (PBMCs) to the *in vitro* DC culture. **(d)** Proportion distributions of pDCs in SLE patients and healthy donors. *FDR* < 0.05. **(e)** Bar plot showing average expression of *PLD4* in pDCs from SLE patients (PBMCs). The Wilcoxon rank sum test was used to measure the statistical significance between SLE patients and healthy controls. **(f)** Schematic of the rs2841277-containing regulatory element with two guide RNA pairs. **(g,h)** Deletion efficiency across amplicon positions for the indicated genes. **(i)** Relative mRNA expression levels of the indicated ISGs in *PLD4*^−/−^x*PLD3*^−/−^ CAL-1 cells rescued with exogenous PLD4. Cells were evaluated under unstimulated conditions or stimulated with RNA9.2s for 4 h (*n* = 5 independent experiments). Statistical analysis was performed using one-way ANOVA followed by Tukey’s post hoc test for multiple group comparisons. **(j)** Lollipop plot of candidate TF regulators of *PLD4* from SCENIC+ colored by triplet score (*z*-score). **(k)** Relative mRNA expression of *TCF4* and *PLD4* after *TCF4* shRNA-mediated knockdown compared with control (shCT) in CAL-1 cells (*n* = 6 independent experiments). Statistical analysis was performed by two-tailed, unpaired Student’s t-test. Data are shown as mean ± s.e.m. \**P* < 0.05; \*\**P* < 0.01; \*\*\**P* < 0.001; \*\*\*\**P* < 0.0001.

